# Hydrophobic Clusters Direct Folding of a Synthetic Chimeric Protein

**DOI:** 10.1101/2023.09.29.560087

**Authors:** Abhishek A. Jalan, Lukas Hassine, Sergio Romero-Romero, Julian Hübner, Kristian Schweimer, Birte Höcker

## Abstract

Extant proteins frequently share sub-domain sized fragments, suggesting that among other mechanisms, proteins evolved new structure and functions via recombination of existing fragments. While the role of protein fragments as evolutionary units is well-established, their biophysical features necessary for generating a well-folded and stable protein are not clearly understood. In order to probe how fragments determine foldability and stability of recombined proteins, we investigated the stability, folding and dynamics of a synthetic chimera created by fusion of fragments of the chemotactic response regulator protein CheY that belongs to the flavodoxin-like fold and imidazole glycerol phosphate synthase from histidine biosynthesis (HisF) which harbors the TIM-barrel fold. The chimera unfolds via an equilibrium intermediate. Mutation of a glycine residue present at the interface of the CheY and HisF fragments to a valine abrogates the equilibrium intermediate while mutation to isoleucine dramatically increases the native state kinetic stability without any significant change in the folding rate. Parts of the fragment interface in the chimera are found to be conformationally dynamic while hydrophobic mutations globally increase its conformational rigidity. We hypothesize that the hydrophobic mutation improves sidechain packing in a large cluster of isoleucine, leucine and valine (ILV) residues that spans the fragment interface. We also extrapolate that inheritance of large ILV clusters from parent proteins could be a key determinant of successful fragment recombination.

## Introduction

Nature frequently refashions new proteins from existing ones using mutation, duplication, recombination, deletion and circular permutation. (1) These genetic events, resulting in emergence of new or diversification of existing protein folds, often leave evolutionary footprints. For example, in-depth sequence and structural analyses have identified several subdomain-sized protein fragments that appear in the context of seemingly unrelated proteins (2–5). Combinatorial shuffling of these fragments is one of the many ways in which proteins evolve (6). While the role of fragments in protein evolution is well-established, the biophysical features of the fragments necessary for generating a foldable and stable protein are not clearly understdood. For example, which fragments lead to a progeny protein that is not only functional but also folds with a reasonable rate and does not misfold or aggregate over its physiological lifetime? Are the fragments intrinsically stable subdomains and thus special enough to be recycled across the proteome? What is the importance of the interface between the fragments for protein foldability and stability? Answers to these questions can improve fundamental understanding of protein evolution and also allow us to define design rules for creating stable proteins with new structures and function.

We can investigate these questions by studying naturally occurring proteins known to have evolved via fragment recombination. However, often fragment-specific information in natural proteins is lost due to mutational drift. For example, the (βα)_8_-barrel topology of the TIM-barrel proteins phosphoribosylformimino-5-aminoimidazole carboxamide ribotide isomerase HisA and the cyclase subunit HisF of the imidazole glycerol phosphate synthase is believed to have arisen from the duplication and fusion of a gene encoding the (βα)_4_ half-barrel (7–10) or even (βα)_2_ quarter-barrel (11). Despite evidence of duplication at the sequence-level, clusters of hydrophobic residues isoleucine, leucine and valine (ILV) known to define cores of stability in HisF are not segregated to the half-barrels (12). Thus, what makes these half-barrels special and how they contribute to HisF folding and stability is difficult to delineate. As an alternative, well-folded and stable synthetic proteins generated using recombination of existing protein fragments can be used to tease out the biophysical features important for successful fragment recombination. We previously developed a synthetic protein chimera containing fragments of the chemotactic response regulator CheY of the flavodoxin-like (βα)_5_-fold (13) and HisF of the TIM-barrel fold. The resulting protein chimera, denoted CheYHisF-1, deviated from the classical (βα)_8_-barrel fold due to the presence of an additional β-strand in the central β-barrel (14). Excision of the residues corresponding to this ninth β-strand resulted in an increased propensity to form higher-order oligomers. Mutation of five residues located at the interface of the CheY and HisF fragments of this construct resulted in the stable and monomeric protein CheYHisF-2, which adopts a TIM-barrel fold (15). The five mutations located at the fragment interface are crucial for the stability and foldability of CheYHisF-2. From an evolutionary perspective, packing of sidechains at the interface between the fragments of a protein newly emerged from recombination is expected to be sub-optimal. Thus, mutations directed at the interface could provide Nature with a tool to remodel the fitness landscape of the recombined proteins. In order to understand the importance of the fragment interface in recombined proteins, we investigated the stability, folding and native state dynamics of CheYHisF-2. Additionally, we also investigated the effect mutations at the fragment interface have on these biophysical features.

We find that CheYHisF-2 is thermodynamically less stable than its topological parent HisF. Additionally, it unfolds via an equilibrium intermediate in the ionic chaotrope GdmCl but no intermediate is observed during equilibrium unfolding in the non-ionic chaotrope urea. Native state dynamics measured using Nuclear Magnetic Resonance (NMR) spectroscopy reveals that parts of the interface between the CheY and HisF fragments are dynamic. Mutation of a single amino acid close to the dynamic region to hydrophobic residues increases the thermodynamic and kinetic stability of the chimera, abrogates the equilibrium intermediate and also increases the stability of the kinetic folding intermediate. This hydrophobic mutation is so located that it can potentially stabilize clusters of hydrophobic ILV amino acids that straddle the fragment interface and extend deep into the CheY and HisF fragments. ILV clusters serve as cores of stability and also play a key role in folding of most proteins. Thus, increased stability of the chimera and the abrogation of the equilibrium intermediate suggests that optimization of hydrophobic clusters across the fragments interface could be a key determinant of successful protein fragment recombination.

## Results and Discussion

### CheYHisF-2 is less stable than the sum of its parts

The thermodynamic stability of CheYHisF-2 was determined via equilibrium chemical unfolding in GdmCl. Denaturation was monitored using change in the fluoresence emission of the single tryptophan (W135) contained in the HisF fragment at 320 nm (excitation = 280 nm) or circular dichroic ellipticity at 225 nm as a function of GdmCl concentration. HisF has been shown to unfold via a two-state process i.e only the native and denatured states are apparent between 0 and 6 M GdmCl (16). The fluorescence and CD unfolding curves of CheYHisF-2 do not overlap (**Fig. 1A**). The fluorescence curve shows a single cooperative transition but the CD curve diverges between 2 and 4 M GdmCl showing an additional unfolding transition. This suggests that significant secondary structure in the chimera persists up to 4 M GdmCl. Thus, unlike HisF, CheYHisF-2 unfolds via an equilibrium folding intermediate. The non-coincidental fluorescence and CD unfolding curves are not due to insufficient equilibration time as the CheYHisF-2 folding is fully reversible (**Fig. S1**). Global fit of the fluorescence and CD unfolding curves in GdmCl to a three-state equilibrium unfolding model (17) yield 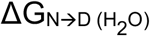 of 30.83 kJ mol^-1^ at 20 ºC (**Table 1**). This value is approximately half of the Gibbs free energy of denaturation of HisF 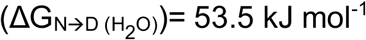 obtained in identical buffer but at 25 ºC (16).

**Table 1.** Thermodynamic parameters of CheYHisF-2 and its mutants. Thermodynamic stability of CheYHisF-2 and its mutants derived from global fitting of the fluorescence and CD unfolding curves to a two-state (urea) or three-state (GdmCl) equilibrium unfolding model. Activation energy (Eact), enthalpy (ΔH) and melting temperature (T_*m*_) were determined by differential scanning calorimetry (DSC).

|  | Chemical Unfolding |  |  |  |  |  | Thermal Unfolding |  |  |
| --- | --- | --- | --- | --- | --- | --- | --- | --- | --- |
|  | GdmCl |  |  |  | Urea |  | DSC |  |  |
| | $\Delta G_{N \rightarrow I}$<br>(kJ mol<sup>-1</sup>) | $\Delta G_{I \rightarrow U}$<br>(kJ mol<sup>-1</sup>) | $m_{N \rightarrow I}$<br>(kJ mol<sup>-1</sup> M<sup>-1</sup>) | $m_{I \rightarrow U}$<br>(kJ mol<sup>-1</sup> M<sup>-1</sup>) | $\Delta G_{N \rightarrow D}$<br>(kJ mol<sup>-1</sup>) | $m_{N \rightarrow D}$<br>(kJ mol<sup>-1</sup> M<sup>-1</sup>) | $E_{act}$<br>(kJ mol<sup>-1</sup>) | $\Delta H$<br>(kJ mol<sup>-1</sup>) | $T_m$<br>(°C) |
| CheYHisF-2 | 19.2 ± 1.0 | 11.7 ± 1.6 | 12.3 ± 0.8 | 4.4 ± 0.4 | 41.7 ± 1.5 | 8.4 ± 0.3 | 86.0 ± 4.5 | 69.9 ± 2.8 | 65.9 ± 0.1 |
| G213A | 14.2 ± 1.0 | 10.0 ± 2.6 | 8.7 ± 0.7 | 4.1 ± 0.6 | 34.0 ± 1.3 | 6.8 ± 0.3 | 65.6 ± 4.2 | 54.7 ± 5.3 | 61.1 ± 0.2 |
| G213V | 22.7 ± 0.8* | - | 12.1 ± 0.4* | - | 44.8 ± 1.8 | 8.5 ± 0.3 | 108.8 ± 7.2 | 79.3 ± 1.4 | 68.9 ± 0.1 |
| G213L | 17.3 ± 1.1 | 13.0 ± 1.8 | 11.3 ± 0.9 | 4.9 ± 0.5 | 47.1 ± 0.9 | 8.5 ± 0.2 | 115.1 ± 9.6 | 64.8 ± 1.1 | 67.3 ± 0.2 |
| G213I | 32.2 ± 3.3 | 12.5 ± 5.1 | 17.2 ± 2.0 | 5.0 ± 1.2 | 50.6 ± 2.4 | 9.1 ± 0.4 | 116.8 ± 9.2 | 84.1 ± 2.1 | 69.4 ± 0.1 |

\*values for N→U transition

**Figure 1.**
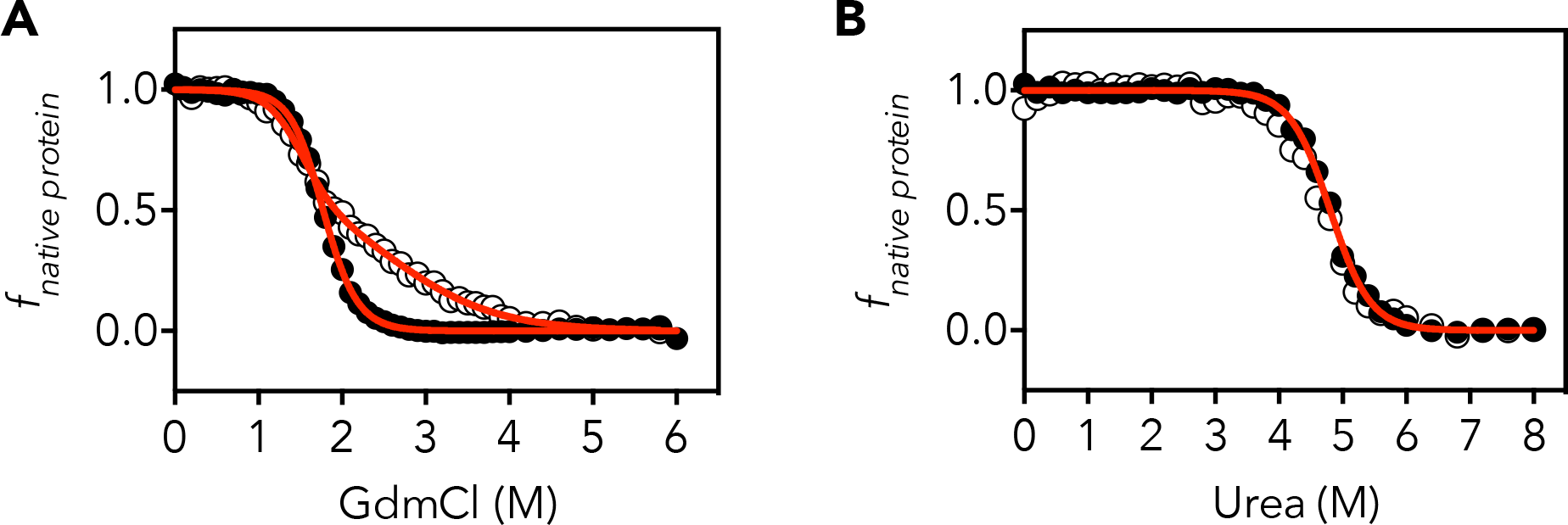
CheYHisF-2 unfolds differently in ionic and non-ionic denaturants. Equilibrium unfolding of CheYHisF-2 obtained in GdmCl (A) and urea (B). Unfolding curves measured via fluorescence emission and CD are shown in solid and open circles respectively. Red lines represent a global fit of the fluorescence and CD unfolding curve to a three-state (GdmCl) or two-state (urea) equilibrium unfolding model.

GdmCl is a monovalent salt and its ionic nature affects the stability of the native state and intermediates via different mechanisms. It can disrupt networks of electrostatic interactions in the native state inducing premature unfolding (18). During unfolding, GdmCl can stabilize near-native (19) as well as non-native (17, 20, 21) clusters of hydrophobic amino acids, populating equilibrium intermediates at moderate denaturant concentrations (22). Given that hydrophobic effect is stronger in ionic solutions (23), we measured equilibrium unfolding of the chimera in the non-ionic chaotrope urea. The CD and fluorescence unfolding curves of CheYHisF-2 in urea overlap (**Fig. 1B**) and the curves are successfully fitted to a two-state equilibrium unfolding model. These results suggest that the equilibrium intermediate observed in GdmCl is populated due to the ionic nature of the denaturant and is likely stabilized by hydrophobic interactions.

Although, CheY and HisF originate from the hyperthermophilic bacterium *Thermotoga maritima*, recombination of their fragments in CheYHisF-2 does not yield a hyperthermophilic protein. Fragment recombination is expected to disrupt native inter-residue interaction at the fragment interface more than the core of the chimera. Previously, a chimeric TIM-barrel created by duplication and fusion of the C-terminal half-barrel of HisF also populated an equilibrium intermediate. The crystal structure of the chimera revealed a long stretch of sequence at the half-barrel interface with missing electron density suggesting disorder in this region. Rational design of the interface recovered the missing electron density, abrogated the equilibrium intermediate and also dramatically increased the chimera’s stability (24). Dynamic regions of proteins have also been previously implicated in stabilizing equilibrium intermediates (25). Is the lower stability of CheYHisF-2 in comparison to HisF and the formation of the equilibrium intermediate a result of unoptimized and hence dynamic fragment interface?

The nine-stranded chimera CheYHisF-1 had been computationally redesigned to obtain CheYHisF-2 with the aid of five mutations. All five mutations were present at the interface of the CheY and HisF fragments. In order to understand the individual contribution of the mutations to the stability of CheYHisF-2, each residue was reverse mutated to its wild-type counterpart. Of the five reverse mutations, G213V most dramatically increased soluble expression level (15). G213 is the only inter-residue contact shared between helix 1 of the HisF fragment and helix 8 of the CheY fragment. Given its position at the fragment interface and the effect on soluble expression, we systematically mutated G213 to either alanine, leucine or isoleucine. Along with G213V, we investigated the equilibrium unfolding of all the mutants in both GdmCl and urea and also determined their thermal and kinetic stability using differential scanning calorimetry (DSC).

### ILV mutations stabilize the chimera and dramatically influence the equilibrium intermediate stability

All the mutants expressed as soluble monomeric proteins as judged by size-exclusion chromatography (SEC) coupled with multiangle light scattering (MALS) shown in **Fig. S2**. Secondary structure analysis by far-UV CD suggested similar proportions of α-helices and β-sheets as well as total secondary structure content (**Fig. S3**). Thermal stability of the CheYHisF-2 and the mutants could not be determined using CD due to aggregation at higher temperatures. In DSC experiments, G213A was thermally least stable while G213I had the highest thermal stability (**Fig. S3C and Table 1**).

The trend in thermal stability was also mirrored by the thermodynamic stabilities of the mutants in both GdmCl and urea (**Fig. 2A and Table 1**) In general, ILV mutations resulted in more stable and compact chimeras as judged by the increase in 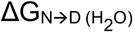 and m-values, respectively. For example, G213A was less stable than CheYHisF-2 by ∼ 7 kJ mol^-1^ in both GdmCl and urea. G213I was more stable by 13.8 kJ mol-1 in GdmCl and 8.9 kJ mol-1 in urea. With the exception of G213A, all the mutants had a higher m-value than CheYHisF-2. Since m-value is a measure of the surface area exposed upon unfolding (26, 27), an increased m-value suggests that the G213I mutant is more compact than CheYHisF-2. If the well-known correlation between ΔASA and m-value is considered, the difference in m-value between G213A and G213I would correspond to a difference of ≅5000 Å^2^ in ΔASA exposed upon unfolding, demonstrating the different degrees of compactness in the studied proteins. This effect is most likely related to the changes in the hydrophobic clusters as discussed below and agrees with the enthalpy changes observed in the temperature-induced unfolding by DSC, where the ΔH value for the G213I mutant is higher than the one for CheYHisF-2 (84.0 vs. 69.9 kcal mol^-1^). When proteins with similar topology and size are compared, the main reasons for finding differences in their unfolding ΔH- and m-values are the number of disrupted internal interactions during the process, as well as the hydration of groups exposed upon unfolding. Both factors correlate with the compactness of the protein. Therefore, chemical- and temperature-induced unfolding experiments demonstrate that the G213I mutant adopts a highly compact conformation in comparison with the other proteins.

**Figure 2.**
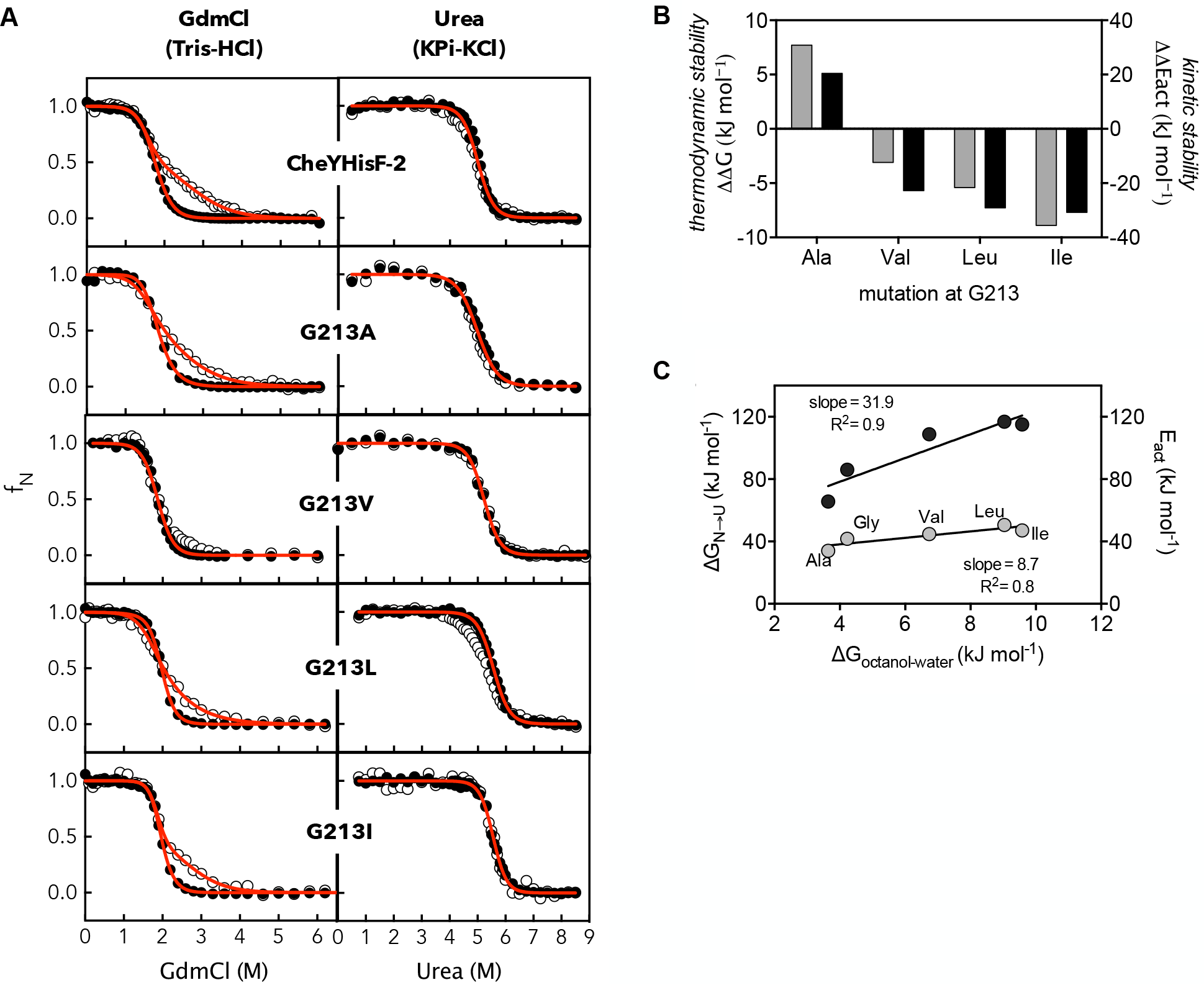
Hydrophobic mutations alter native state and equilibrium intermediate stability. **(A)** Equilibrium unfolding curves of CheYHisF-2 and the mutants obtained in GdmCl and urea. Open and closed circles represent fluorescence and circular dichroism unfolding data. Red lines denote the fit of fluorescence and CD unfolding curves to a two and a three-state equilibrium unfolding model for urea and GdmCl, respectively. (**B**) Change in thermodynamic (ΔΔG in grey bars) and kinetic (ΔΔE_act_ in black bars) stability of the chimera upon mutation of G213 to alanine, valine, leucine or isoleucine. (**C**) Linear-regression analysis and correlation of the thermodynamic (ΔG_N→U_ in grey circles) and kinetic (E_act_ in black circles) stability to Wimley-White octanol-water transfer free-energies (28) of the hydrophobic side chains. R-squared Pearson correlation coefficients and slope of the linear regression fit are indicated. The thermodynamic stability of all data points is reported from the chemical unfolding in urea.

DSC traces of CheYHisF-2 and its mutants showed an increase in T_*m*_ by varying the scan-rate and no endotherm was obtained after a reheating cycle, indicating that their thermal unfolding is under kinetic control (**Fig. S4**). Since their thermal unfolding follows an irreversible two-state mechanism (N → F), it was possible to determine the activation energy (E_act_) between the native and the transition state, which is a measure of the kinetic stability. For all proteins, coincident values are obtained from the fitting of the endotherms to the model and as well for the slope from the Arrhenius plot (**Fig. S4**). When E_act_ is compared among mutants, kinetic stability is ranked progressively from G213A → CheYHisF-2 → G213V → G213L → G213I, in agreement with the increase observed in the thermodynamic stability by chemical-induced unfolding. Thus, G213I is both thermodynamically as well as kinetically more stable than CheYHisF-2 (**Fig. 2B**).

Previous discussion about the linkage between thermodynamic and kinetic stability has suggested that there are two fundamental scenarios, which depend on the functional protein being thermodynamically stable or not with respect to the final states (unfolded or partially unfolded) (29). In the case of CheYHisF-2 and its mutants, since their thermal unfolding is well-described by the Lumry–Eyring model (**Fig. S6**), the native state is thermodynamically stable (as indicated by the chemical-unfolding experiments). But this state undergoes irreversible alteration processes, leading to a final state that is unable to fold back to the native state. According to this scenario, the experimental results indicate that both thermodynamic and kinetic stabilities are coupled, thus the mutations introduced in CheYHisF-2 show parallel effects on both parameters, i.e. G213A has the lowest ΔG and E_act_ values, while the G213I mutant exhibits the highest ΔG and E_act_ values. In addition, a consequence of having very different activation energy values is that the kinetic stability also varies over a broad range. This is reflected in extreme changes in half-life times (t_1/2_) among mutants, e.g. from 275 days for G213A to 2.65 x 10^5^ years for G213I.

Hydrophobic interactions within an ILV cluster can modulate the stability and folding of proteins via non-specific hydrophobic interactions as well as sidechain packing density (30). Glycine and alanine mutations result in under-packed ILV clusters. This could explain the lower stability of G213 and G213A in comparison to G213V, G213L and G213I, which provide non-specific Van der Waals hydrophobic interactions and also induce higher packing density within the ILV cluster. The degree of thermodynamic as well as kinetic stability of CheYHisF-2 correlates with the Wimley-White octanol-water transfer free energies (28) of the mutated hydrophobic sidechains (**Fig. 2C**). The transfer energies are a measure of non-specific hydrophobic interactions. Thus, the correlated increase in stability with increase in hydrophobicity of the mutated sidechains is not surprising. However, the effect of the differences in the packing density due to isoleucine, valine or leucine mutations is not captured by stability alone. It has been shown that the environment of hydrophobic residues is evolutionarily optimized for better packing density and that even a switch between isoleucine and valine, which differ only in a single methyl group, can result in a substantial change in thioredoxin stability (30). In our case, while we do see a moderate change in stability between G213V, G213L and G213I, the effect of difference in the packing density is better captured by the stability of the equilibrium intermediate observed during chemical unfolding in GdmCl. Of the five mutant constructs, only G213V does not populate an equilibrium intermediate during chemical unfolding. It is likely that glycine and alanine result in an under-packed ILV cluster while leucine and isoleucine, both of which contain an extra methyl group compared to valine, overpack the cluster.

Wu *et al*. have previously proposed that tightly packed clusters of ILV residues guide the folding of TIM-barrel proteins and by extrapolation (βα)n parallel-strand proteins. For example, selective mutation of residues in the ILV cluster of the alpha subunit of tryptophan synthase to alanine destabilizes a kinetic folding intermediate (31). Given that hydrophobic mutations at G213 strongly influences equilibrium intermediate stability, we investigated if these also affect the folding pathway and the stability of kinetic folding intermediates. We studied the unfolding and refolding kinetics of CheYHisF-2 and G213I in urea. These two variants were chosen because they sit on the opposite ends of the trend in thermodynamic and kinetic stability.

### Hydrophobic mutation stabilizes the kinetic intermediate of CheYHisF-2

In order to comparatively assess the stability of the kinetic folding intermediates of CheYHisF-2 and G213I, we first set out to unravel their folding mechanisms. Unfolding and refolding kinetics of CheYHisF-2 and G213I were measured by intrinsic tryptophan fluorescence emission at 320 nm and CD ellipticity and 225 nm. In a typical experiment, 40 μM native protein unfolded in 8M denaturant was manually diluted 10-fold to varying final concentrations of urea and the observed time traces were fit to a suitable sum of exponential functions. Unfolding and refolding of both variants followed a mono-exponential and bi-exponential kinetic, respectively. The total amplitude of the fluorescence and CD unfolding kinetic traces was similar to the difference between the native and unfolded signals obtained from equilibrium unfolding measurements (**Fig. S5**). In contrast, kinetic refolding traces of both variants showed ∼40% loss in fluorescence amplitude within the deadtime of the experiment at 1.0 M final urea concentration (**Fig. S6A**). The magnitude of amplitude loss and the refolding rates remained unchanged with a ten-fold drop in the final protein concentration from 4 to 0.4 μM for CheYHisF-2 (**Fig. S6B**) suggesting that the loss in amplitude was not due to aggregation. The initial value of the refolding traces obtained by extrapolation to time zero decreased linearly with an increase in denaturant concentration for both variants (**Fig. S7A**). While this could have suggested reversible aggregation, the initial and final values of the unfolded and refolding kinetic traces obtained via stopped-flow mixing accounted for the total change in fluorescence amplitude (**Fig. S7B**) without any change in the relaxation rates (**Fig. S7C**). This suggests that the missing amplitude during manual mixing is likely an experimental artefact and not due to a fast folding species, generally referred to as the burst phase, populated in the deadtime of the refolding experiment. The relaxation time (∼10s) of the fast refolding phase is close to the 8-10s deadtime of manual mixing. This could cause errors in extrapolation of the refolding traces to time zero resulting in the observed experimental effect.

The unfolding rate constant of both variants monitored by CD overlapped with those obtained using fluorescence but the refolding rate constants show partial overlap (**Fig. S8A**). The fast phase rates observed by CD show large dispersion around those observed via fluorescence while the slow phase rates overlap beyond 3.0 M urea. Intriguingly, the rate constants obtained by a mono-exponential fit of the refolding traces overlap with the slow phase obtained by fluorescence (**Fig. S8B**). In all likelihood, this is a result of the intrinsic low sensitivity of the CD instrument and the inability to measure the decay of the fast phase accurately. An increase in concentration to 8 or 12 μM protein did not improve the dispersion in the rate constants of the fast phase. This trend was observed for both the CheYHisF-2 and the G213I variants. Thus, we relied on the rate constants and amplitudes obtained from the mono-exponential fit of the refolding traces to perform burst-phase analysis.

Kinetic refolding monitored by CD showed a ∼70% loss in amplitude at 1.0 M urea during the deadtime of the experiment for both variants (**Fig. S9**). The loss in amplitude was independent of protein concentration but, unlike fluorescence, the initial value of refolding kinetic traces obtained by extrapolation to zero time decreased in a sigmoidal fashion with increasing denaturant concentration (**Fig. 3A**). The sigmoidal trend and the magnitude of amplitude loss during deadtime was independent of protein concentration. Thus, the observed change likely did not arise from reversible aggregation. This suggests that the polypeptide of both variants rapidly collapses with ultrafast kinetics into a burst-phase intermediate (I_BP_). Given the strong similarity of the kinetic folding profile of our chimeras to those reported for HisF as well as synthetic chimeras sym1 and sym2 (24), it is unlikely that the folding kinetics of the burst-phase would fall in the time regime measurable by stopped-flow mixing.

**Figure 3.**
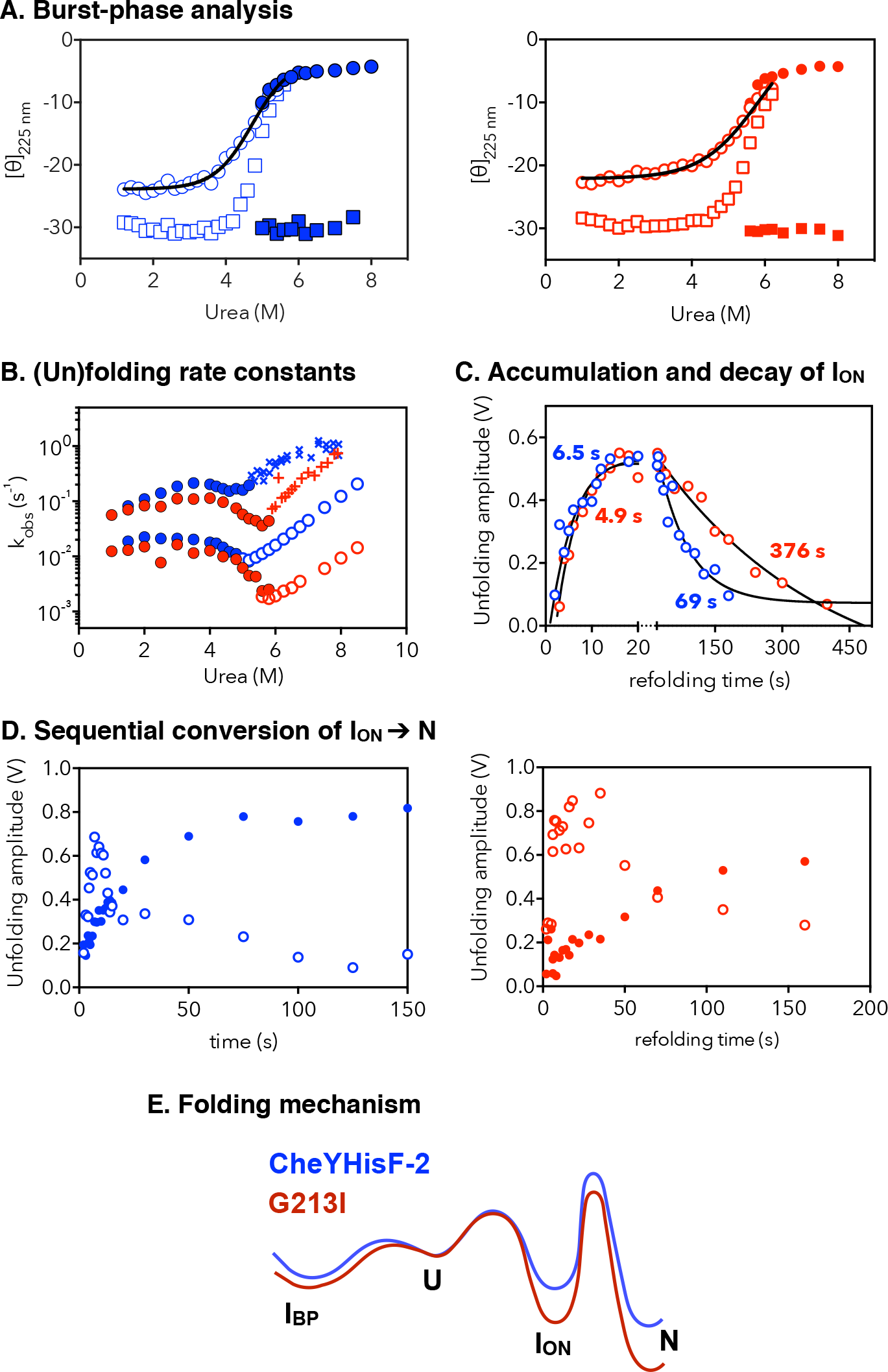
Folding kinetics of CheYHisF-2 (blue) and G213I (red). (**A**) Burst-phase analysis of the amplitudes of unfolding and refolding kinetic traces obtained by monitoring the CD signal at 225 nm. The initial (circle) and final (square) values of the refolding (open symbols) and unfolding (closed symbols) are shown along with a two-state fit (solid line) of the initial value of refolding. (**B**) Refolding (filled symbols) and unfolding (open symbols) rate constants obtained using stopped-flow mixing with excitation at 280 nm and emission > 320 nm. The crossed symbols represent I_ON_ unfolding rates obtained by refolding the proteins in 4.0 M urea for 18 s and then unfolding between 5.18 and 7.93 M urea concentration. (**C**) Unfolding amplitudes obtained by refolding the proteins in 4.0 M urea followed by variable time delay and then unfolding in 5.18 M urea (CheYHisF-2) or 5.9 M urea (G213I). The solid lines represent fit of the unfolding amplitudes to a biexponential-equation and the numbers show the obtained rate constants. (**D**) Unfolding amplitudes obtained by refolding the proteins in 4.0 M urea for a variable period followed by unfolding in 8.5 M urea. The filled and empty circles represent unfolding amplitudes of the native state and the on-pathway intermediate, respectively. (**E**) Schematic illustration of the folding mechanism depicting burst-phase intermediate (I_BP_), unfolded state (U), on-pathway intermediate (I_ON_) and the native state (N) of CheYHisF-2 and G213I.

Fit of the burst-phase curves to the two-state equilibrium unfolding model resulted in a ΔG_D_ of 22.0 kJ mol^-1^ and m-value of 4.7 kJ mol^-1^ M^-1^ for CheYHisF-2 and a ΔG_D_ of 18.1 kJ mol^-1^ and m-value of 3.1 kJ for G213I, suggesting that the I_BP_ of CheYHisF-2 is moderately more stable and compact than the one of G213I. It is remarkable that both HisF and CheYHisF-2 share the propensity to form a burst-phase intermediate despite the substitution of an 80 amino acid fragment from the N-terminal part of HisF with the structurally similar fragments from CheY. It has been proposed that the information for early-folding kinetic intermediates is encoded into the primary sequence of the proteins. This raises the intriguing possibility that the burst-phase intermediate observed in HisF could be localized to the C-terminal half-barrel. A plot of the observed unfolding and refolding relaxation rates versus urea concentration resulted in a chevron plot (**Fig. 3B**) that bears strong similarity to the one for HisF (16). The unfolding rates of both variants decreased linearly with an increase in urea concentration. In contrast, the refolding rates of the fast phase showed a pronounced positive slope until ∼ 4.0 M urea followed by a decrease in rates until an inflexion point at ∼ 4.8 M urea. The rates increased again beyond the inflexion point resulting in the canonical chevron shape of the fast phase. The initial positive slope of the fast phase suggests that the molecular species it represents unfolds faster as urea concentration increases. This is consistent with the formation of a folding intermediate. The refolding rates observed in the slow phase are independent of denaturant concentration until ∼ 4.0 M for both variants followed by a decrease until the inflexion point. The denaturant-independent folding kinetics, also generally referred to as “roll-over” in the context of chevron plots, suggests the formation of an intermediate in the slow phase as well. The amplitude of the fast phase increases while that of the slow phase decreases until 3.0 M beyond which the trend is reversed. Taken together, the observed roll-over and the reciprocal relationship in their folding amplitude suggest that the two phases represent two sequentially formed intermediates on the folding pathway of both the variants. Similar to HisF, the fast phase likely represents the conversion of the I_BP_ into another intermediate I_ON_ and the slow phase its subsequent conversion into the native state. The subscript “ON” indicates the on-pathway nature of the intermediate represented by the slow-phase.

Interestingly, while the onset of the negative slope in the fast phase of both variants is similar at ∼ 4.0 M, the inflexion point for the G213I mutant is higher by 0.8 M urea. Additionally, its amplitude also persists until a higher urea concentration as compared to the fast phase amplitude of CheYHisF-2. These observations suggest that the folding intermediate of G213I is more stable than that of CheYHisF-2. This is also supported by the large difference in the unfolding rate of the intermediate as determined by stopped-flow double-jump experiments presented below. The slope of the slow refolding phase of both variants is similar but the inflexion point of the G213I mutant is higher than CheYHisF-2 by 0.8 M urea. This combined with the difference in the unfolding rate of the slow phase accounts for the increased thermodynamic stability of the G213I mutant.

Given the rapid interconversion of the intermediates with the unfolded polypeptide, we could not determine whether the I_BP_ is an on- or an off-pathway intermediate. The I_BPs_ observed in HisF and the two synthetic chimeras sym1 and sym2 generated from duplication and fusion of its half-barrel have previously been shown to be an off-pathway intermediate that unfolds before progressing to the on-pathway intermediate via a low kinetic barrier. Therefore, it is reasonable to assume that the I_BP_ of both variants in our case is also an off-pathway intermediate.

We further investigated the likely formation of I_ON_ in the fast phase and its sequential decay in the slow phase using stopped-flow interrupted refolding assays. In order to first verify if the I_ON_ is indeed formed in the fast phase, unfolded protein was refolded in 4.0 M urea for 18 s and the refolding interrupted by transferring to 7.9 M urea. At 4.0 M urea, the time constant of the fast and the slow phase are 5 and 55 s for CheYHisF-2 and 9 and 78 s for G213I, respectively. Thus, during the refolding episode lasting 18 s, a major fraction of the polypeptide population should refold into the intermediate. Concomitantly, its unfolding during the second episode should be faster than that expected of the native proteins under similar conditions. Indeed, the molecular species populated after the second jump decays with a time constant of 0.4 s for CheYHisF-2 and 0.7 s for G213I (**Fig. S10**), which are respectively one and three orders of magnitude higher than the time constants observed for the unfolding of these proteins in single mixing experiments. These results suggest that the fast refolding phase represents the formation of I_ON._

Using an identical experimental setup, we determined the unfolding limb of the I_ON_ by transferring the protein solutions refolded for 18 s to varying concentrations of urea between 5.18 and 7.93 M. The rates obtained are plotted against urea concentration in **Fig. 3B**. The unfolding limb of I_ON_ obtained by the interrupted-refolding experiments joins the refolding limb obtained in single mixing experiments and result in a chevron shape for both variants. This suggests that the folding of I_ON_ is reversible.

The temporal evolution and decay of I_ON_ were investigated using further interrupted-refolding experiments, which follow the I_ON_ amplitude as it changes with increase in the refolding time. This can be used to compare the relative stability of the I_ON_ in both variants. Unfolded CheYHisF-2 or G213I were allowed to refold in 4.0 M urea and after variable time delay of 1-500s transferred to 5.18 or 5.9 M urea, respectively. These urea concentrations fall in the refolding limb of the slow phase but in the unfolding limb of the fast phase. As a result, I_ON_ formed in the fast phase unfolds but the native state formed in the slow phase remains folded. As expected, the rate constant of I_ON_ does not vary while the amplitude first increases, reaches a plateau and then decreases with time delay (**Fig. 3C**). Curiously, a biexponential fit of the unfolding amplitude reveals that the I_ON_ of CheYHisF-2 and G213I accumulates with time constants of 6.5 and 4.9 s but decays with time constants of 69 and 376 s. These time constants match well with those obtained in single mixing experiments, suggesting that they indeed represent the accumulation of I_ON_ in the fast phase and its decay to the native state in the slow phase, respectively. In additional interrupted refolding experiments, we investigated if this process is sequential.

In a sequential reaction, the amplitude of the species populating second would always show a lag in comparison to the first one. The large difference in the unfolding rates of the I_ON_ and the native states of both variants (**Fig. 3D**) ensured that we could simultaneously follow their unfolding amplitude by refolding the polypeptides for variable delay and then unfolding in 8.5 M urea. A clear lag in the unfolding amplitude of the native state of both variants is observed. The evidences presented above suggest that the folding mechanism of CheYHisF-2 and G213I as shown in **Fig. 3E** is conserved vis-à-vis HisF.

### Hydrophobic mutation increases the conformational rigidity of CheYHisF-2

Mutation of G213 to leucine, valine or isoleucine stabilizes the chimera, strongly influences stability of the equilibrium and kinetic intermediates and also renders the protein more compact. Such a multifaceted change in the biophysical properties of the chimera is likely a result of improved hydrophobic packing in G213I. This would restrict the backbone and sidechain dynamics in the mutant protein. Motion within chimera can occur on multiple time scales. We focused on two that relate most closely to the stability and folding of the chimera; internal friction on the nanosecond to millisecond timescale and conformational fluctuation on the minutes to hour timescale. The fast dynamics provides clues to suboptimal packing at the fragment interface as well as global compactness of the proteins while the slower dynamics measures how tightly the different secondary structure elements are packed within the tertiary structure. The fast and slow timescale motions in CheYHisF2 and G213I were measured using longitudinal (T1) and transverse (T2) relaxation and hydrogen-deuterium exchange (HDX) NMR spectroscopy.

The backbone ^15^N-amide relaxation parameters of most amides in CheYHisF-2 are characteristic of a protein of this size (**Fig. 4**). Importantly, the relaxation rates of the CheY and HisF fragments are similar. Based on the very uniform R_2_/R_1_ ratios the overall molecular tumbling can be described very well by an isotropic tumbling with a rotational correlation time of 11.1 ns for CheYHisF-2 and 10.4 ns for G213I at 313K. For G213I R_1_ is slightly higher (about 5%) and R_2_ lower (about 5%) compared to CheYHisF-2 resulting in the reduced rorational correlation time. This might suggest an enhanced compactness supporting the its higher m-value of G213I obtained via chemical unfolding in urea, but translational diffusion experiments with the same samples resulted in an identical diffusion coefficient (1.33 10^-10^ m^2^/s for CheYHisF-2 and 1.33 10^-10^ m^2^/s for G213I) suggesting no difference of the hydrodynamic radius. The highly uniform R_2_/R_1_ ratios suggest that the two fragments form a cohesive single-domain protein and tumble as a single unit. Two regions show enhanced R_2_/R_1_ ratios due to elevated R_2_ rates as a result of chemical exchange on the μs-ms timescale. The loop connecting strand β6 and helix α6 (residues 153-160) undergoes chemical exchange on the μs-ms timescale and is spatially close to the phosphate binding site observed in HisF. The chemical exchange could be a result of weaker binding to phosphate in CheYHisF-2 as compared to HisF resulting in significant populations of bound and unbound chimera. The loop connecting strand β8 and helix α8 (residues 209-212) also shows chemical exchange on a the μs-ms timescale. This loop is located close to the fragment interface. Importantly, residue position 213 is positioned at the end of this loop. In the G213I variant the contribution of chemical exchange to R_2_ is reduced.

**Figure 4.**
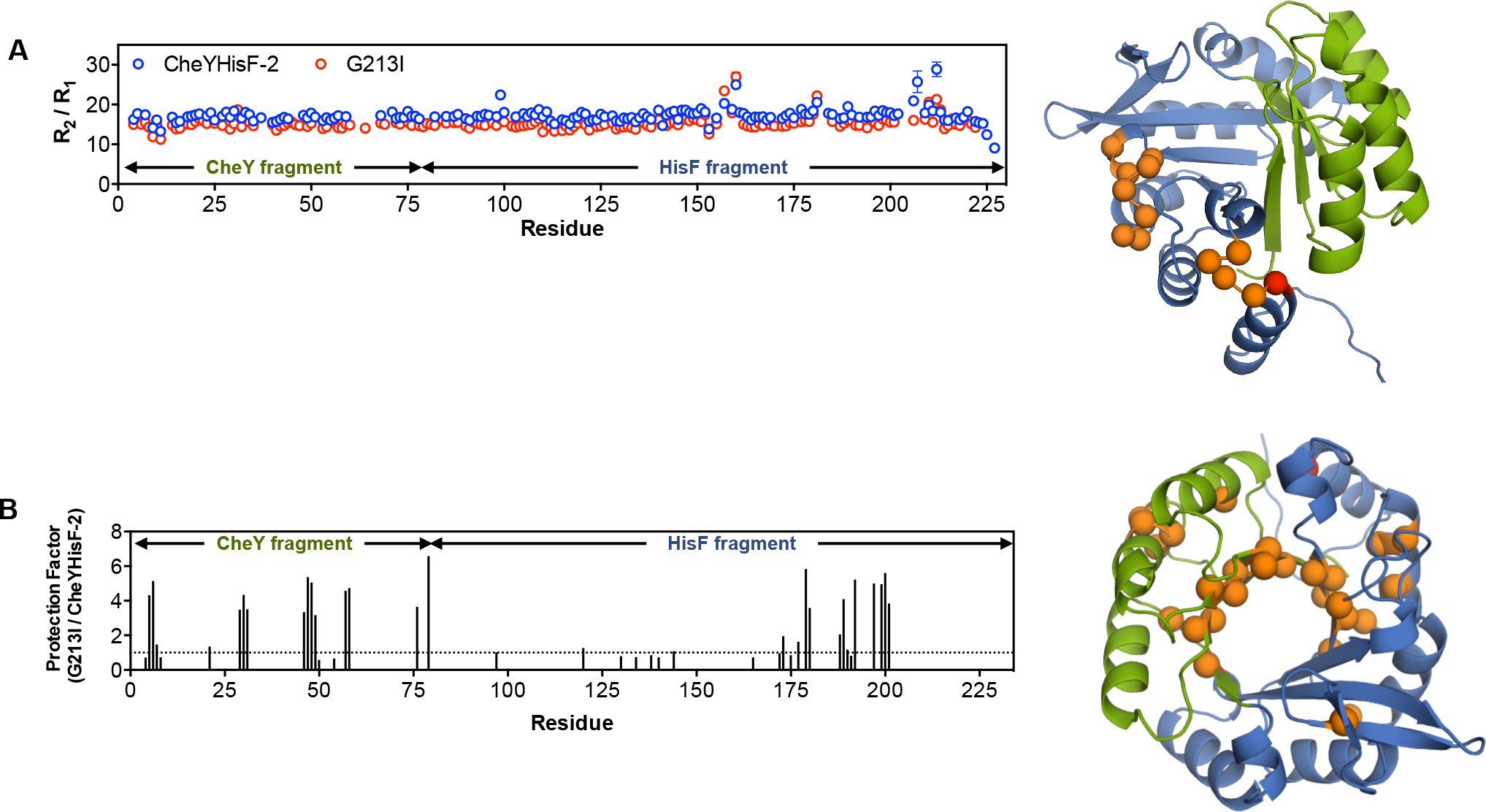
Effect of hydrophobic mutation on the conformational dynamics of CheYHisF-2. **A**) Backbone dynamics represented as the ratio of transverse (R2) and longitudinal (R1) relaxation rates for CheYHisF-2 (blue) and G213I (red). Cα atoms of residues that show chemical exchange on the μs-ms timescale are highlighted as orange spheres. (**B**) Ratio of protection factors of G213I and CheYHisF-2 obtained from hydrogen-deuterium exchange NMR spectroscopy. Cα atoms of residues that show a protection factor ratio > 1 are mapped onto the CheYHisF-2 structure as orange spheres. The CheY and HisF fragments are shown in green and blue, respectively, and the location of residue 213 is highlighted in red.

We also measured the effect of G213I mutation on the ability of the backbone amides to exchange with the solvent using hydrogen-deuterium exchange (HDX) spectroscopy. As previously observed in the case of HisF (12), three classes of amides are apparent; class I amides that exchange within the time required to dissolve the lyophilized sample in D_2_O, shim the magnet and acquire the first spectra (∼15 minutes), class II amides that decay to > 90% of the initial intensity and class III amides that decay to less than 90% of the initial intensity. The decay in intensity of class II amides were fit to a single exponential function and the rate constants k_ex_ normalized with respect to intrinsic exchange rate K_int_ calculated based on a random coil polypeptide. The resulting values, called protection factors, measure resistance of amides protons to exchange with deuterons. As shown in **Fig. 4B**, the majority of class II amides in G213I have a higher protection factor than CheYHisF-2. More importantly, the amides with increased protection are distributed in both the CheY and HisF fragment. This indicates that the mutation globally restricts the conformational dynamics rather than only in the vicinity of the mutated residue.

### Hydrophobic interactions across the interface could be important for successful fragment recombination

Hydrophobic interactions guide folding of nearly all globular proteins. In the specific case of HisF, ILV sidechains serve as cores of stability and determine a protein’s stability and folding (12, 18). These clusters also stabilize partially folded high energy states in other TIM-barrel proteins (31). In our case, mutation of G213 to ILV residues strongly influences the stability of the chimera and the equilibrium intermediate. As observed in double-mixing stopped flow experiments, the amplitude of the I_ON_ of CheYHisF-2 decays at a faster rate than that of G213I, suggesting that mutation to isoleucine increases the stability of the on-pathway folding intermediate. Natural proteins have evolved to conserve ILV clusters within related proteins if not always at the same location in sequence (12). This bears special significance in case of fragment recombination given the progeny protein may not inherit intact clusters from the parent proteins. How does partial or complete inheritance of ILV clusters influences the foldability and stability of the chimera? Could the merger of the inherited clusters into large continuous clusters that span the fragment interface be a prerequisite for emergence of well-folded proteins? Here, we assess these questions by analysing the ILV clusters in crystal structures of CheY and HisF and the chimera CheYHisF-2.

ILV clusters are self-contained hydrophobic units within a folded or partially folded protein that together bury more than 500 Å^2^ total sidechain surface area and at least 10 Å^2^ area between two contacting residues (32). The webserver BASiC (31) and the Contacts of Structural Units (CSU) software (33) that were previously used to identify ILV clusters in proteins are no longer available online. Thus, we reimplemented the original CSU algorithm recently to enable identification of ILV clusters in the webserver ProteinTools (34). CheY from *Thermotoga maritima* contains two large ILV clusters burying 1038 (cluster 1) and 1868 Å^2^ (cluster 2) surface area and contain 44 and 23 inter-residue contacts, respectively (**Fig. 5**). Residues in cluster 1 are primarily contributed by α_4_ and α_5_, both of which are excised during fragment recombination and not inherited in the chimera. Cluster 2 is formed by residues of the α_2_(αβ)_3_β_4_ element, which is transferred nearly intact into the chimera. HisF contains four large hydrophobic clusters burying 888 (cluster 2), 1041 (cluster 3), 1503 (cluster 4) and 3758 Å^2^ (cluster 5) surface area. Cluster 3 formed by residues of the (βα)_1−2_β_3_ element is completely removed during recombination. Cluster 5 containing 96 inter-residue contacts in the α_3−_β_7_ element is only partially retained in the chimera. This creates a region at the fragment interface between helices α_4_ and α_5_ of the chimera that is completely devoid of any hydrophobic clusters. Clusters 0 and 4 of HisF are inherited nearly entirely by the chimera. More importantly, these two clusters merge with cluster 2 of CheY to create a large continuous cluster that straddles the fragment interface between helices α_1_ and α_8_. The cluster that emerged from this merger contains 39 inter-residue contacts and buries 1750 Å^2^ surface area. To summarize, one cluster from CheY and two from HisF merge to create two large clusters in the chimera, of which only one spans the fragment interface. Residue position 213 which so dramatically affects the stability of the native state as well as its equilibrium and kinetic intermediates is located within this large cluster spaning the interface.

**Figure 5.**
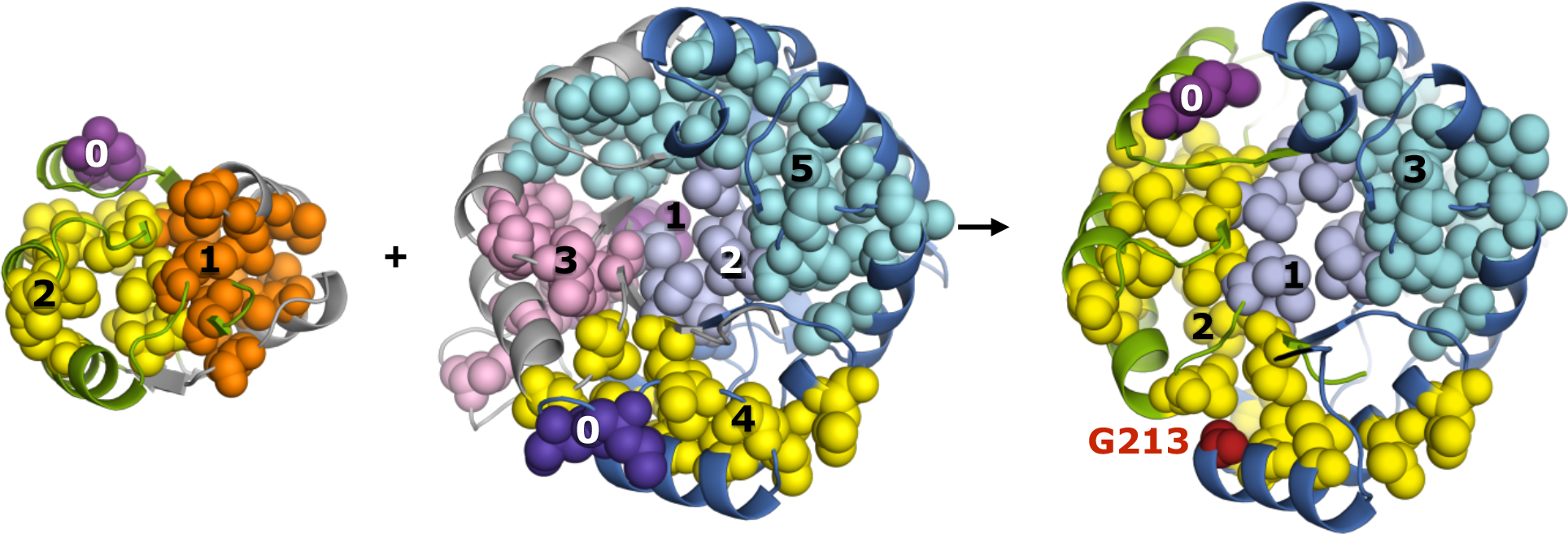
Distribution of hydrophobic clusters in CheY, HisF and CheYHisF-2. CheY (left) and HisF (middle) fragments retained in CheYHisF-2 (right) are shown in green and blue, respectively, and the parts removed during recombination are shown in grey. Clusters are numbered in decreasing order of the number of inter-residue contacts; for example, cluster 0 has the lowest and cluster 3 has the highest number of inter-residue contacts in CheYHisF-2. The position of G213 in cluster 2 of the chimera is shown in red.

### Implications of mutations at fragment interfaces

Mutation is a singularly important tool in the arsenal of protein evolution. Nature selects for mutations that slow the unfolding rate i.e. increase the kinetic stability. (35, 36) While the folding rate also increases over an evolutionary time scale, a positive or negative correlation between the residue conservation and folding speed is not observed (37). Thus, nature seemingly selects more strongly for decreased unfolding speed than increased folding speed. The increased selection for kinetic stability protects the protein against misfolding and aggregation (29) while also offering greater evolvability via generous acceptance of deleterious mutations (38). As a protein born from fragment recombination traverses the mutational landscape, mutations in which regions would have a greater chance of increasing kinetic stability? It is established that different parts of a protein evolve at different rates. Amino acid sites critical to function are under purifying selection and evolve slowly compared to the non-functional regions. The rate of site-specific substitutions also correlates with the sidechain packing density (39). Densely packed regions evolve slowly because mutations that alter the packing density stress the system by destabilizing the active conformation and thus are considered deleterious (40). In the specific case of fragment recombination, a sizable fraction of the natively packed protein core is inherited from the parent proteins but even in the best-case scenario, where the choice of crossover point minimizes loss of native contacts, non-native interactions are expected to abound at the interface. These interfacial residues, although present in the protein core, would have lower sidechain packing density and likely evolve faster than the core sites inherited intact from parental proteins. Thus, mutations at the fragment interface would have a greater chance of increasing kinetic stability of the recombined proteins.

It is instructive to note that when the nine-stranded CheYHisF-1 was redesigned into CheYHisF-2 all five mutations suggested by the Rosetta software were present at the fragment interfaces. These mutations, while stabilizing the eight-stranded TIM-barrel, also increased the soluble expression level and general stability of the chimera. Rosetta’s fixed backbone protein design protocol used for the chimera redesign iteratively mutates and samples rotamers to improve sidechain packing and increase protein stability. Computational accumulation of all five mutations at the interface suggests that indeed protein fragment recombination benefits from improved sidechain packing at the interface. As observed with CheYHisF-2, the suboptimal packing also results in a dynamic interface, which can more easily tolerate small amplitude rigid-body movements needed to remould the interfacial packing density through mutations. In light of these observations, it can be argued that within the limit of stability, suboptimal packing at the fragment interface promotes protein evolvability due to greater opportunity for sculpting residue-level interactions. From the perspective of chimeragenesis-based protein design, mutations at the interface will likely be less deleterious and rational or computational approaches to improve chimera stability might be focused in these regions.

### Potential of synthetic chimeras in determining biophysical constraints on fragment recombination

As discussed in the previous section, fragment interfaces in recombined proteins are likely to accumulate mutations at a faster rate than the rest of the protein. Such mutations will improve the sidechain packing at the interface and change the global stability and dynamics of the protein. Thus, any fragment-specific information encoded in the sequence that drove successful recombination would likely be lost to time on an evolutionary scale. Consequently, biophysical features that make the frequently recycled protein fragments special and constraint their recombination into new proteins are difficult to elucidate using natural proteins. Synthetic chimeras are an excellent alternative primarily because mutational perturbations at the fragment interface can provide clues to biophysical constraints on fragment recombination. Furthermore, Nature only contains examples of successful fragment recombinations. However, synthetic chimeragenesis is not always successful and can result in aggregation, misfolding, formation of an unintended protein fold and so on. These diverse outcomes can be rationalized at the sequence level leading to a better understanding of the factors influencing successful chimeragenesis.

For example, fusion of fragments from the TIM barrel HisF and the chemotactic response regulator CheY of the flavodoxin-like fold results in a chimera CheYHisF-1 (**Fig. 1**)(14) with a fold similar to TIM-barrels but with an additional ninth β-strand in the central barrel. Removal of the strand-forming residues and computational redesign resulted in a second chimera CheYHisF-2 with a proper TIM-barrel fold.(15) We have previously created two additional chimeras NarLHisF and MMCoAHisF created from the fusion of HisF with fragments from *Escherichia coli* nitrite response regulator NarL and methylmalonyl CoA mutase (MMCoA) from *Propionibacterium shermanii*, respectively. NarLHisF is stable and monomeric and modelling suggests a TIM-barrel like fold but MMCoAHisF on the other hand overexpresses insolubly and aggregates upon refolding (41). Analogous to fragment recombination in Nature, CheYHisF-1 and NarLHisF represent evolution of a new fold, CheYHisF-2 represents sequence diversification and MMCoAHisF represents a failed recombination.

The molecular basis of such different outcomes of fragment recombination can be investigated using synthetic chimeras via two complementary approaches. The first is primarily a design-based approach where a significant library of topologically related chimeras such as CheYHisF, NarLHisF, MMCoAHisF etc. is created by mining structurally similar proteins of the flavodoxin-like fold at the organismal and superfamily level and then experimentally validated. Trends in thermodynamic stability, soluble recombinant expression levels and resistance to proteolytic degradation can then be empirically rationalized based on relative contact order, solvent accessible surface area, distribution of hydrophobic cluster and other known biophysical determinants of protein stability. In order for the rationalization to be statistically significant, a large library of soluble, monomeric and stable chimeras must be generated, by no means a trivial task. The second approach is to study the folding and fragment-specific dynamics of the available successful chimeras to understand the molecular basis of their stability. We have followed this later approach and have elucidated the folding and dynamics of the TIM-barrel like CheYHisF-2 using equilibrium and kinetic measurements and NMR spectroscopy. These have revealed that sidechain packing at the interface and continuity of hydrophobic clusters in the fragments is important for successful recombination. In future, investigations into folding and dynamics of CheYHisF-1 and NarLHisF as well as computational and rational approaches to rescuing MMCoAHisF will further shed light on the molecular basis of successful fragment recombination.

## Conclusion

Recombination of protein fragments is a plausible mechanism for emergence of new and diversification of existing proteins. Recent efforts have identified several fragments within the context of seemingly unrelated protein folds, suggesting conservation at the subdomain level. (42, 43) Knowledge of these subdomain level evolutionary units is incomplete without understanding the biophysical features that constrain their recombination into stable and functional proteins. The role of specific fragments in determining foldability and stability of recombined natural proteins is expected to be mutationally remodelled over an evolutionary timescale. This makes it difficult to determine the contribution of fragments and the inter-fragment interface to the stability and folding of natural proteins. To circumvent this problem, we investigated the folding and dynamics of a synthetic protein CheYHisF-2 obtained through recombination of fragments from two different protein folds. This chimeric protein is built from fragments of hyperthermophilic proteins but its thermodynamic stability is significantly lower than its topological parent protein HisF. Mutation of residue position 213 located at the fragment interface to isoleucine, leucine or valine increases the thermodynamic and kinetic stability of the chimera. The increase in stability is correlated to the hydrophobicity of the mutated residues. Furthermore, the chimera unfolds via an equilibrium intermediate in GdmCl and its stability is also extremely sensitive to the type of mutation. Measurement of the backbone dynamics using NMR spectroscopy reveals that the hydrophobic mutation G213I renders the chimera more compact and conformationally rigid. Clearly, such a multifaceted change in the biophysical properties of the chimera is a result of improved hydrophobic packing in the protein core owing to mutation. In light of these observations, we extrapolate that the optimization of hydrophobic contacts across the fragment interface is a likely determinant of successful fragment recombination.

## Materials and Methods

### Heterologous Expression and Purification of Proteins

The five CheYHisF-2 variants, which all carry a His_6_-tag at their C termini, were expressed in *E. coli* BL21(DE3). The cells were grown at 37°C in Luria broth supplemented with 100 μg/mL ampicillin for plasmid maintenance. At an OD_600_ of ∼ 0.6 expression as induced by adding isopropyl-β-thiogalactoside (IPTG) to a final concentration of 1mM. Growth was allowed for another 18 h at 20 °C. Cells were harvested by centrifugation (Beckman Coulter, JLA 8.1000, 15 min, 4000 x g, 4 °C) and washed with 50 mM KPi containing 300 mM KCl at pH 7.5 and centrifuged again (Eppendorf centrifuge 5810R, 60 min, 2800 x g. 4 °C). The five proteins were mainly found in the soluble fraction of the cell extract and purified from there. The cells were resuspended in 20 mL resuspension buffer (50 mM KPi, 300 mM KCl and 20 mM imidazole pH 7.5) and lysed by sonification (Branson 6.3 mm tip, 3 x 2 min, Output 4, 40% pulse, on ice) and the resulting homogenate was centrifuged (Beckman Coulter, JA 25.50, 40.000 x g, 60 min, 4 °C). The supernatant was filtered and loaded onto a HisTrap Hp NiNTA column (GE, Life Science, Marlborough) preequilibrated with resuspension buffer. The protein was eluted using a linear gradient of resuspension buffer containing 500 mM imidazole. Fractions with the highest protein content were pooled, loaded onto a Superdex 75 HiLoad 26/60 column (GE, Life Science, Marlborough) and isocratically eluted with 50 mM KPi and 300 mM KCl at pH 7.5. Protein concentrations were determined using the molar extinction coefficient of 11460 M^-1^ cm^-1^ at 280 nm calculated from the amino acid sequence.

### SEC-MALS

SEC-MALS was performed using a Superdex 75 Increase 10/300 GL column (GE, Life Sciece, Marlborough) attached to an ÄKTA Pure sytem coupled to a miniDAWN multi-angle light scattering detector with a laser (658.7 nm wavelength) and an Optilab refractometer (Wyatt Technology, Snata Barbara, CA). All experiments were performed at room temperature (25 °C). Data collection and SEC-MALS analysis were performed with ASTRA 7.3.2 software (Wyatt Technology). The refractive index of the solvent was defined as 1.3309 and the viscosity was defined as 0.8902 with dn/dc values of 0.1850 ml/g. The proteins were eluted with a speed of 0.8 mL/min in 50 mM KPO4 300 mM KCl pH 7.5 with 0.02 % sodium azide.

### Thermal melts followed by Circular Dichroism

Measurements to determine the far-UV CD spectrum and the melting courve were performed using a JASCO model J815 CD spectrophotometer (path-length, 2 mm; bandwidth, 1 nm) with protein solutions of 4 μM concentration (50 mM KPO4 300 mM KCl pH 7.5). CD spectra were recorded from 200 – 250 nm in 10 repetitions and for he Tmelt a wavelength of 225 nm was chosen. The melting courve was recorded with an increasing temperature gradient (0.5 °C/min). Measurements to determine the fluorescence spectra were performed using a JASCO model FP-6500 spectrofluorometer. Protein tertiary structure was analyzed with a spectra from 300 to 400 nm emission scan (bandwidth 5 nm) after excitation at 280 nm (bandwidth 3 nm) for all variants. T-melts were obtained monitoring CD signal at 225 nm as a function of temperature using a protein concentration of 4 μM mg mL^-1^ in buffer 50 mM KPi 300 mM KCl pH 7.5, a heating rate of 1.0 °C min^−1^, and a 2 mm cuvette with a bandwith of 1 nm. CD signal was normalized by:

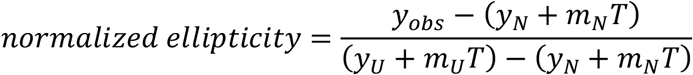

where y_obs_ is the experimentally ellipticity signal at a given temperature, and (*y*_*N*_+*m*_*N*_T) and (*y*_*U*_+*m*_*U*_T) are the linear fits corresponding to the native and unfolded transitions, respectively. *T*_*m*_ values were estimated from normalized data fitted with a Boltzmann-type function:

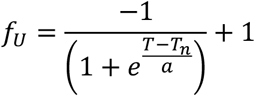

### Thermal unfolding experiments followed by Differential Scanning Calorimetry

Differential Scanning Calorimetry (DSC) endotherms were collected in a VP-Capillary DSC system (MicroCal®, Malvern Panalytical). Samples at 0.5 mg mL^-1^ were prepared by exhaustive dialysis in buffer 50 mM KPi and 300 mM KCl pH 7.5 and then degassed at room temperature. In all cases, proper instrument equilibration was performed by running two buffer-buffer scans before each protein-buffer experiment, and corresponding buffer-buffer traces were subtracted from each protein-buffer scan. For all proteins, irreversibility in the thermal unfolding was observed after a reheating cycle. Endotherms were collected varying scan rate from 1.0 to 3.0 °C min^-1^. Calorimetric transitions were well-fitted by the two-state irreversible model (N→F), where the kinetic conversion from native protein (N) to the final state (F) is described by a first-order constant (*k*) changing with the temperature according to the Arrhenius equation (1):

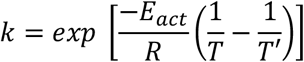

where *T’* is the temperature at which the *k* = 1 min^−1^ and *E*_*act*_ is the activation energy that describes the thermal unfolding process between the native and the transition states. The apparent heat capacity (*C*_P_ ^APP^) is given by:

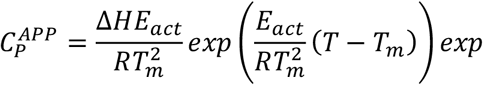

where *T* and Δ*H* are the temperature at a specific value and the unfolding enthalpy, respectively. The *E*_*act*_ was also calculated from the slope of Arrhenius plots, i.e. ln *k* vs. 1/T. Origin v.7.0 (OriginLab Corporation, Northampton, MA, USA.) with MicroCal software was used for data analysis.

### Equilibrium and Kinetic Unfolding and Refolding

GdmCl and Urea mediated equilibrium (un)folding of CheYHisF-2 as shown in Fig. 1 were performed in 50 mM tris-HCl at pH 7.5 to match the experimental conditions reported for HisF. However, some of the mutants took longer to equilibrate in urea due to its weaker denaturation capability. During this delay, the mutants containing no or low concentrations of urea started showing signs of degradation as judged from the progressive loss in their CD signal. Thus, chemical unfolding in GdmCl was measured in tris-HCl buffer at pH 7.5 but unfolding in urea was measured in 50 mM potassium phosphate (KPi) and 300 mM KCl at pH 7.5. The fluorescence measurements to analyze the changes in tertiary structure were carried out with a JASCO-6500 spectrofluorometer. Protein (un)folding induced by GdmCl or urea was followed by the decrease of the fluorescence signal at 320 nm (bandwidth 5 nm) for all variants after excitation at 280 nm (bandwidth 3 nm). The change in secondary structure was followed by far-UV circular dichroism (CD) signal at 225 nm using a JASCO model J815 CD spectrophotometer (path-length, 2 mm; bandwidth, 1 nm). The proteins were incubated with different concentrations of the denaturant and the signals were recorded after different time intervals until no further change was observed. Temperature-induced unfolding was analyzed by following the far-UV CD signal at 225 nm at a scan rate of 1 ºC min^-1^.

For rapid kinetics, fluorescence emission was followed using a 320 nm cutoff filter (cGMP) after excitation at 280 nm in a stopped-flow SX17 Spectrometer from Applied Photophysics (Leatherhead, UK). The samples reached equilibrium within the first 3 days at 25 °C. The transitions were analyzed using a two- or three-state equilibrium model. A linear dependency of the free-energy of unfolding and refolding on the denaturant concentration was assumed. The unfolding and refolding rates were determined by fitting monoexponential or bi-exponential equations to the data points using the software GraphPad PRISM 6. The unfolding limbs of the intermediates were determined by stopped-flow interrupted refolding experiments (double jump; see main text).

### NMR Spectroscopy

Data were recorded on a Bruker AVANCE IIIHD 700, 900 and 1000 MHz spectrometers equipped with ^1^H/^15^N/^13^C cryo-probes. For each measurement, a 1 mM protein sample containing 50 mM potassium phosphate, 300 mM KCl at pH 7.5 was diluted with 10 % v/v D_2_O and 160 μL sample poured into a 3 mM NMR tube. BEST-TROSY-based triple resonance data (2) were recorded for resonance assignment of the G213I variant using non-linear sampling (NUS) with 25% of total data points. Data processing was performed with in-house software (K.S., unpublished) based on the iterative soft thresholding method (3); NMRViewJ (OneMoon Scientific) was used for visualization and analysis. Backbone amide resonance assignments for WT were taken from published data (4). For characterization of overall and internal motions, ^15^N longitudinal (*R*_1_) and transverse (*R*_2_) relaxation rates at 700.2 MHz ^1^H frequency at a calibrated temperature of 313 K. Relaxation delays of *R*_1_ and R_2_ relaxation experiments were fitted to a mono-exponential decay using NMRViewJ (OneMoon Scientific). Rotational correlation time was determined based on the R_2_/R_1_ ration using the program tensor2 (5).

Translational diffusion was measured at 700 MHz ^1^H frequency. Gradient strength was calibrated using a doped water sample (2mM CuSO_4_, 1% H_2_O in D_2_O) assuming a diffusion coefficient of 1.90 * 10^-9^ m2/s at a calibrated temperature of 298 K using a 1D ^1^H pulse gradient stimulated echo with bipolar gradients (6). For protein measurements a calibrated temperature of 313 K was used and the 3-9-19 watergate water suppression was applied in the diffusion experiment. The integrated signal intensity of the methyl group region (0.5 – 1.25 ppm) was observed as a function of increased gradient strength, and analysed according to S(Q) = S(0) * exp{-DQ} with Q = γ^2^g^2^δ^2^(Δ-δ/3-τ/2) (γ = gyromatic ratio, g = gradient strength, δ = gradient pulse length, Δ = diffusion delay, τ = gradient recovery delay (7). For each sample four independent measurements were performed and the corresponding intensitites averaged before data fitting.

For the amide proton-deuterium exchange, 1 mM protein samples containing 50 mM potassium phosphate, 300 mM KCl at pH 7.5 were frozen in liquid nitrogen and lyophilized overnight. The lyophilized protein samples were resuspended in 99.98 % D_2_O, immediately transferred to the probe and a series of 1H, 15N HSQC experiments recorded at 313 K. The time between dissolution of the protein sample and recording of the first HSQC spectra was ∼ 15 mins. The HSQC cross peaks were fit to a monoexponential function to obtain the decay rates *k*_*obs*_. Residue-specific protection factors (PF) were calculated as the ratio of the observed decay rate to the intrinsic exchange rate *k*_*int*_ expected of the same residues in a random coil polypeptide. The *k*_*int*_ were obtained from the online webserver Sphere (www.fccc.edu/research/labs/roder/sphere/) using default settings except at pH of 7.5 and temperature 313 K.

## Supplementary Figures

**Figure S1.**
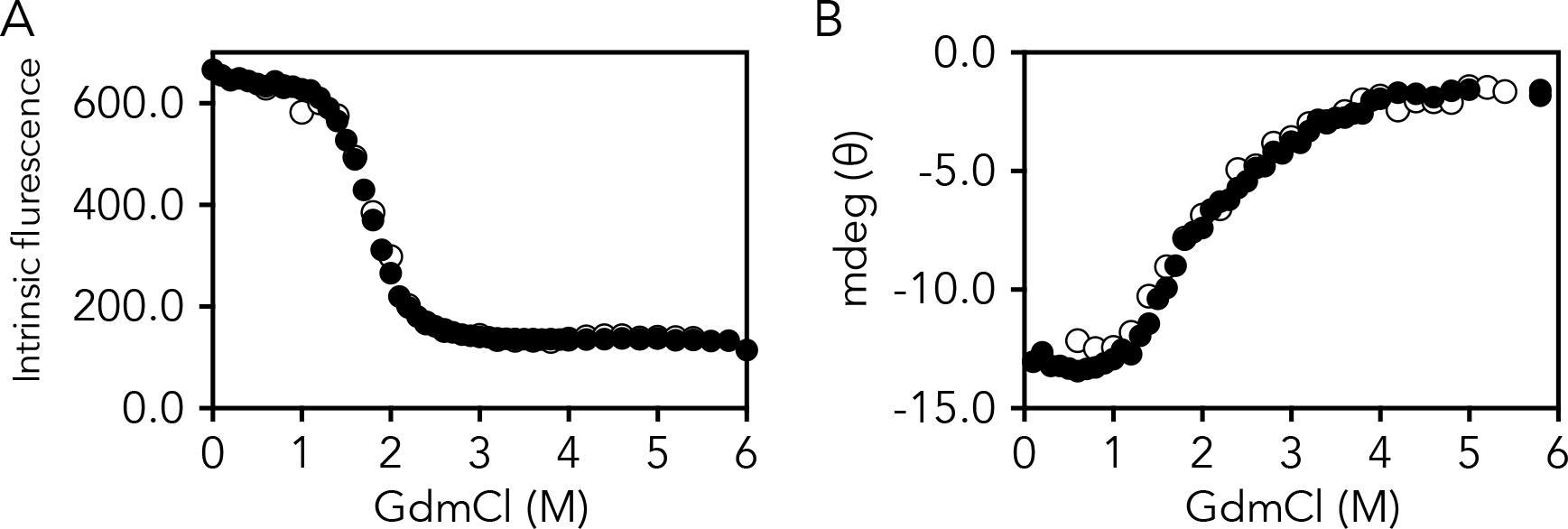
Reversibility of chemical unfolding in GdmCl. Chemical unfolding (closed circles) and refolding (open circles) of CheYHisF-2 monitored using fluorescence (left, excitation = 280 nm, emission = 320 nm) and CD (ellipticity at 225 nm). All experiments were performed at 20 ºC in the presence of tris-HCl buffer at pH 7.5 and 4 μM protein concentration.

**Figure S2.**
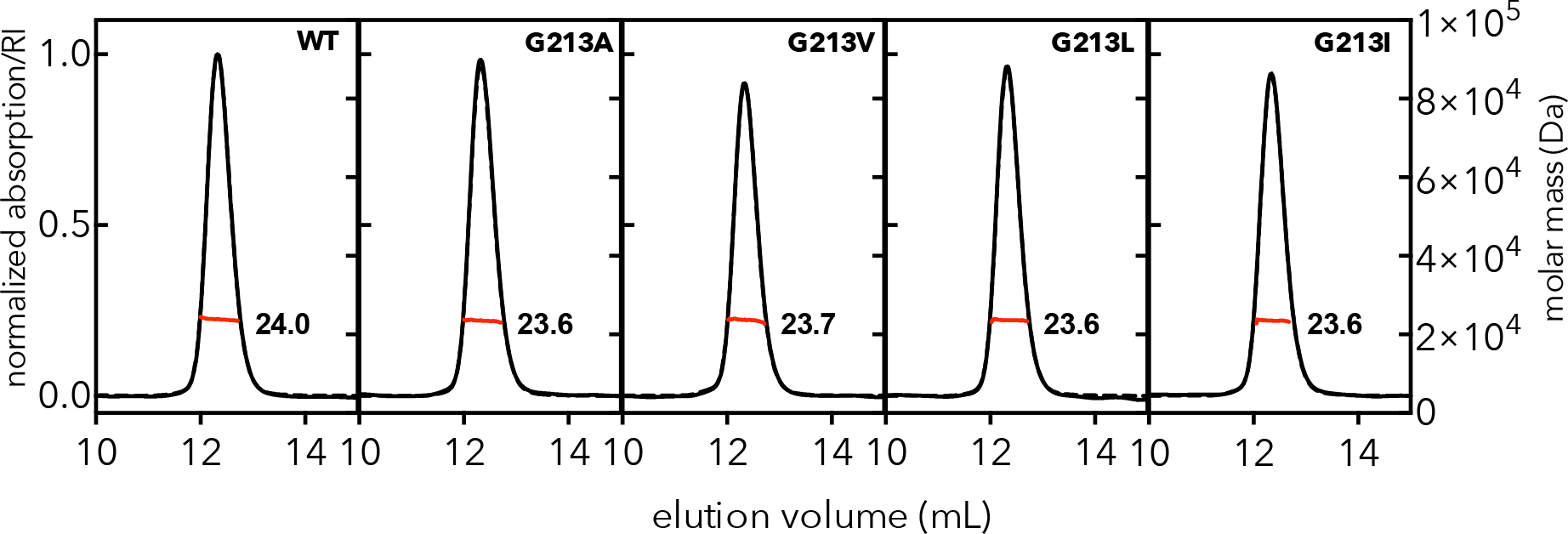
Size-exclusion chromatography coupled multiangle light scattering (SEC-MALS) chromatograms of CheYHisF-2 and it five mutants. The normalized differential refractive index (solid lines), UV absorption at 280 nm (dashed lines) and MW determination (red line) are indicated. The numbers represent molar mass M_*W*_ in units of kDa.

**Figure S3.**
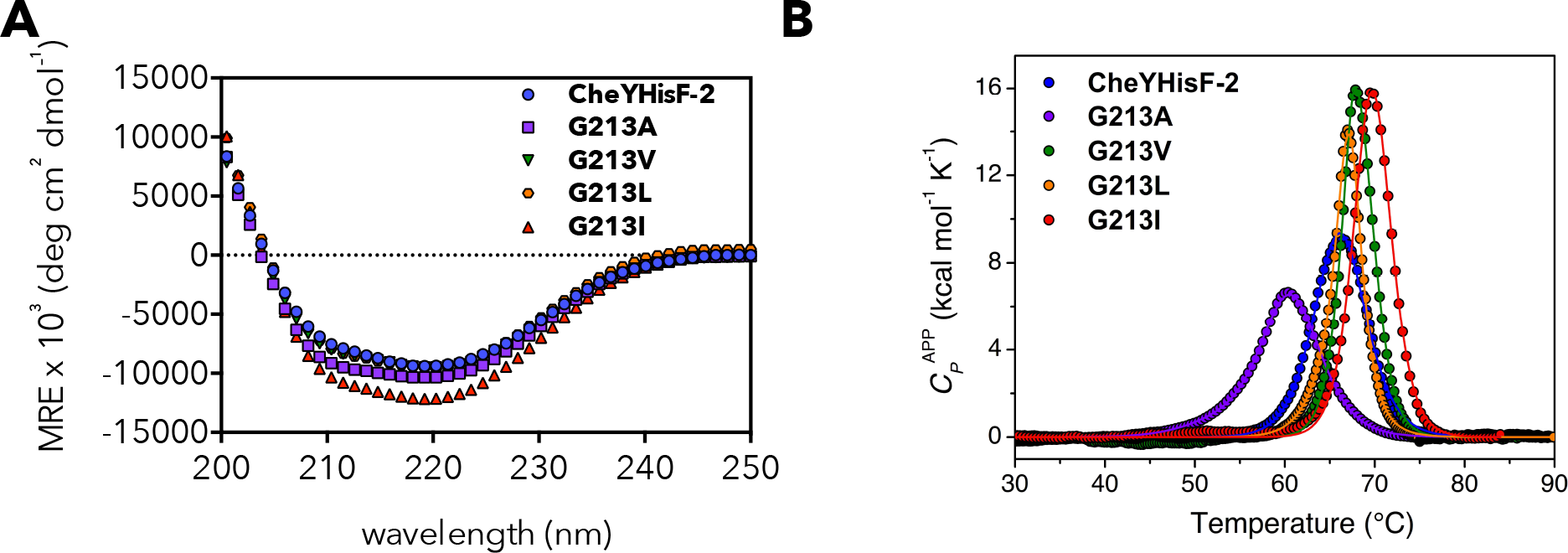
Far-UV circular dichroism spectrographs (**A**) and thermal melts obtained by DSC (**B**) of CheYHisF-2 and its mutants at 4 μM protein concentration in presence of 50 mM potassium phosphate buffer and 300 mM KCl at pH 7.5.

**Figure S4.**
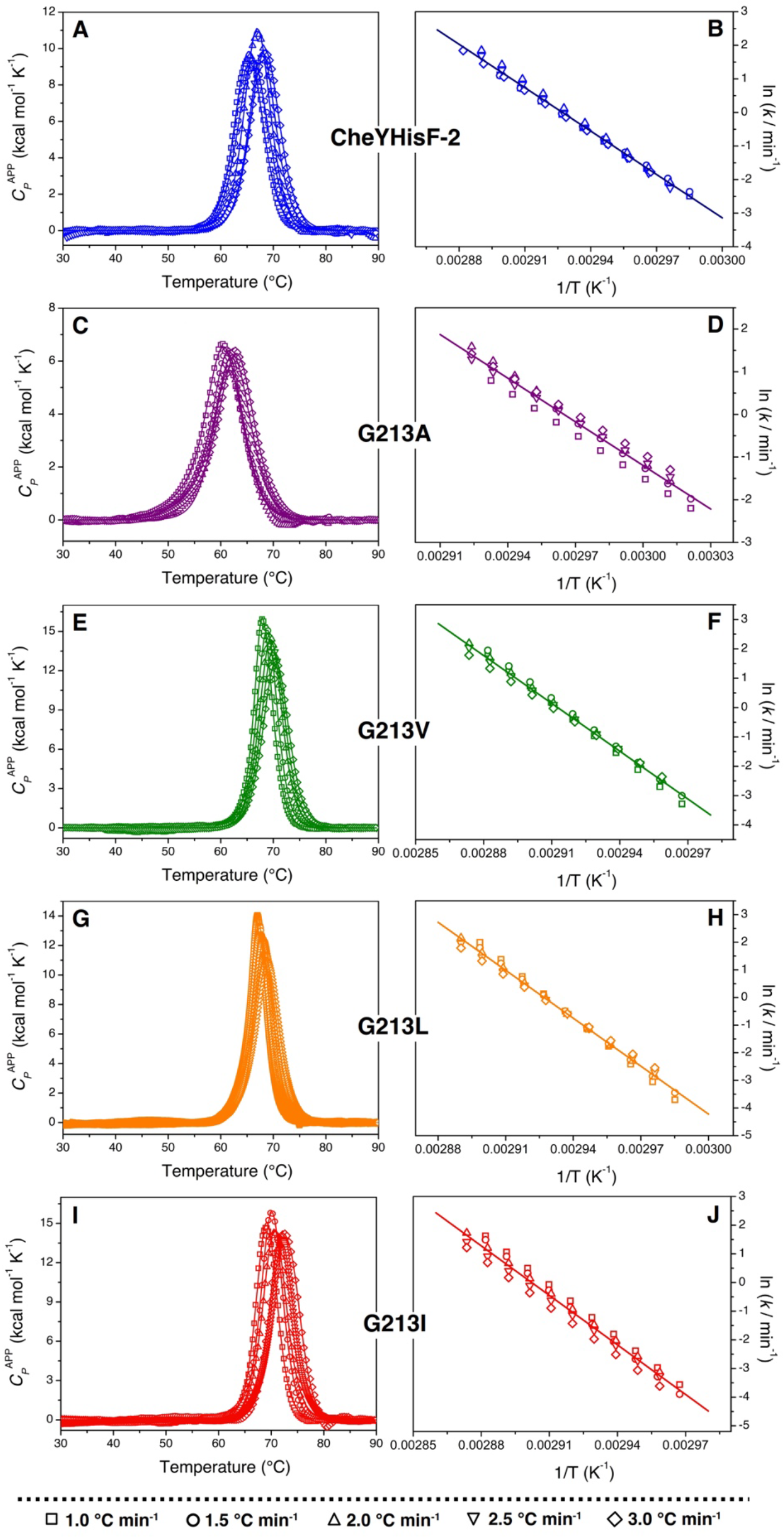
Temperature-induced unfolding of CheYHisF-2 and its mutants. Endotherms at 0.5 mg mL^-1^ and different scan rates (1, 1.5, 2, 2.5, and 3 °C min^-1^) are shown in panels A, C, E, G, and I; continuous lines represent the best fit to a two-state irreversible model (1). Arrhenius plots also including data at different scan rates are presented in panels B, D, F, H, and J; the line shows the best fit to the Arrhenius equation. Protein concentration 0.5 mg/ml in buffer 50 mM KPi and 300 mM KCl pH 7.5.

**Figure S5.**
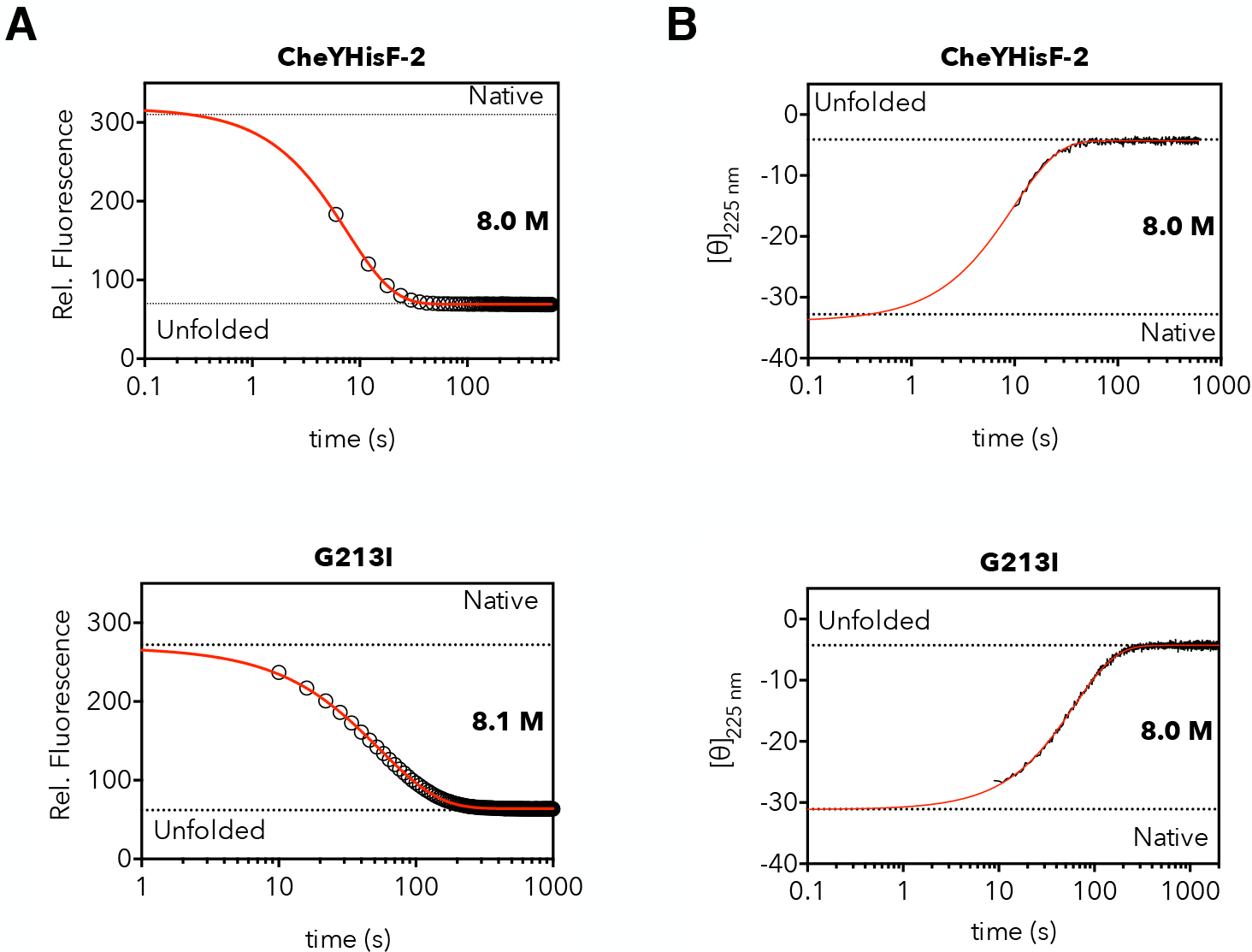
Kinetic unfolding time traces of CheYHisF-2 and G213I as obtained by fluorescence (**A**) and CD (**B**) at 4 μM protein concentration. The dashed lines represent the signals of native or unfolded protein obtained from the fit of pre- and post-transition region of the equilibrium unfolding experiments discussed in the main text. Fit of the traces to an equation describing monoexponential decay is shown in red.

**Figure S6.**
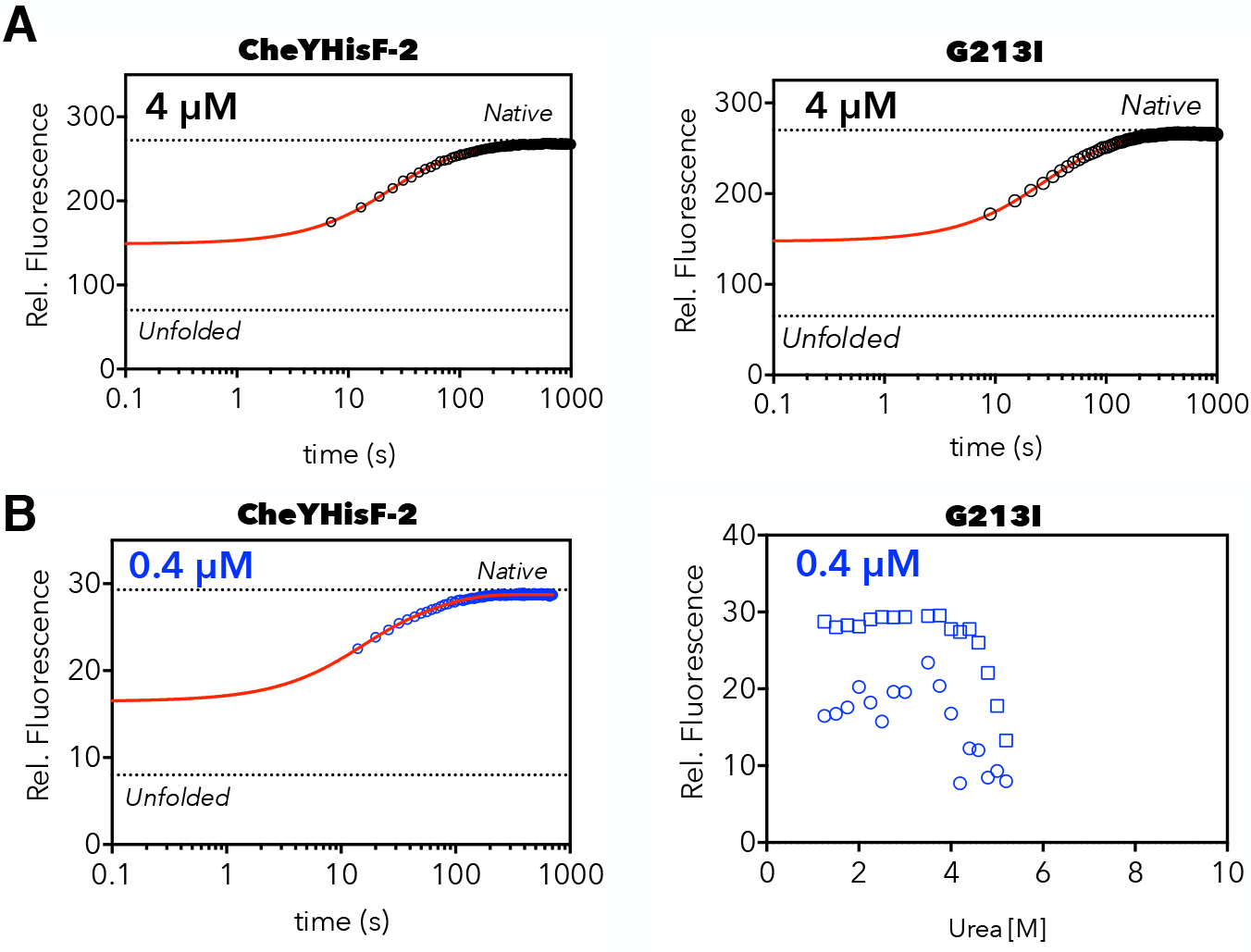
Kinetic refolding traces of CheYHisF-2 and G213I at 4 μM (A) and of CheYHisF-2 at 0.4 μM (blue) protein concentration. All refolding traces were recorded at 1.0 M final urea concentration. Fit of the traces to an equation describing biexponential decay is shown in red.

**Figure S7.**
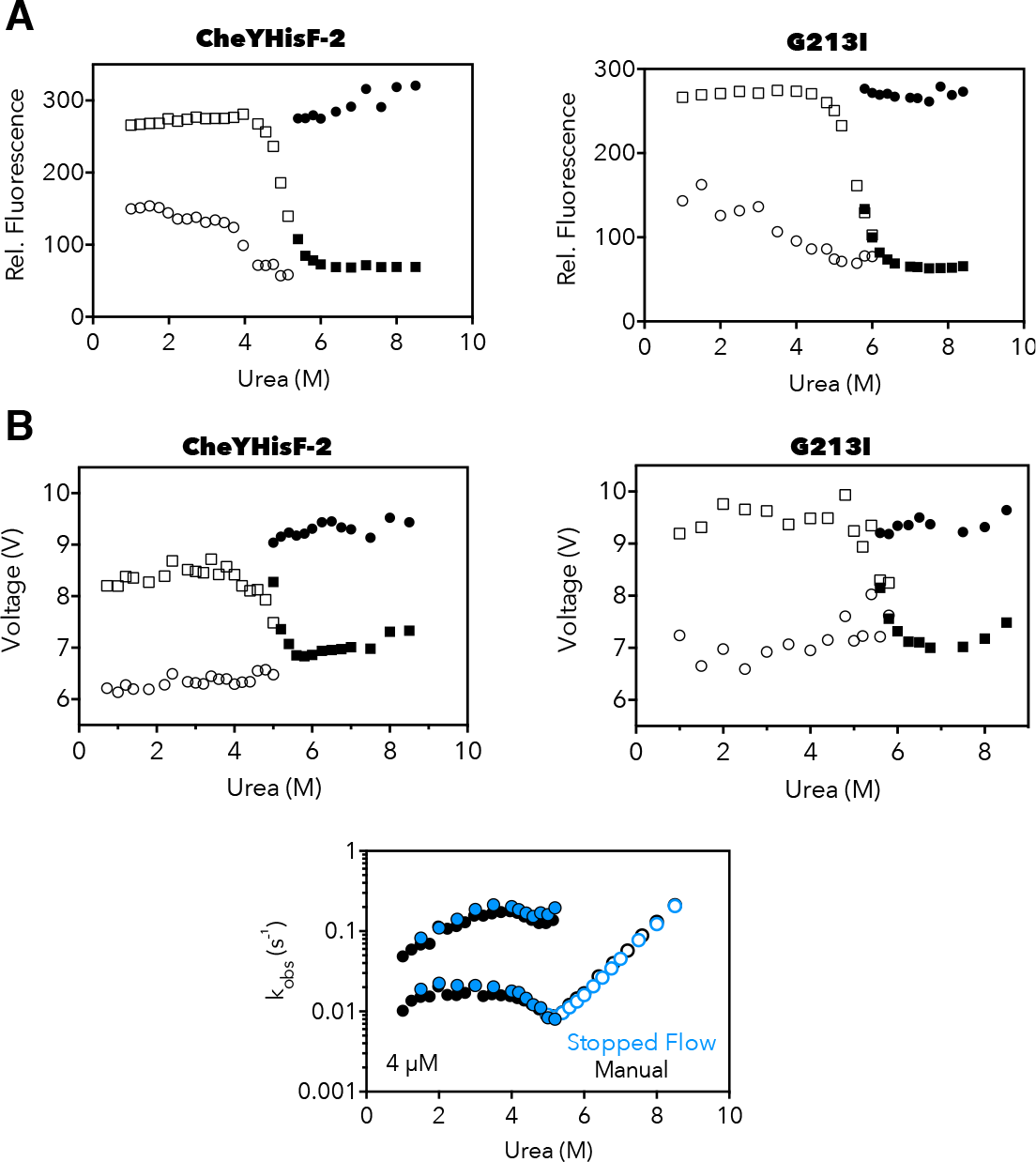
Initial (circles) and final (square) values of refolding (open symbols) and unfolding (closed symbols) of CheYHisF-2 and G213I obtained by extrapolation of fluorescence kinetic traces recorded using manual (**A**) and stopped-flow mixing (**B**) to time zero. The chevron plot of CheYHisF-2 obtained using manual and stopped-flow mixing. All experiments were performed at 4.0 μM protein concentration.

**Figure S8.**
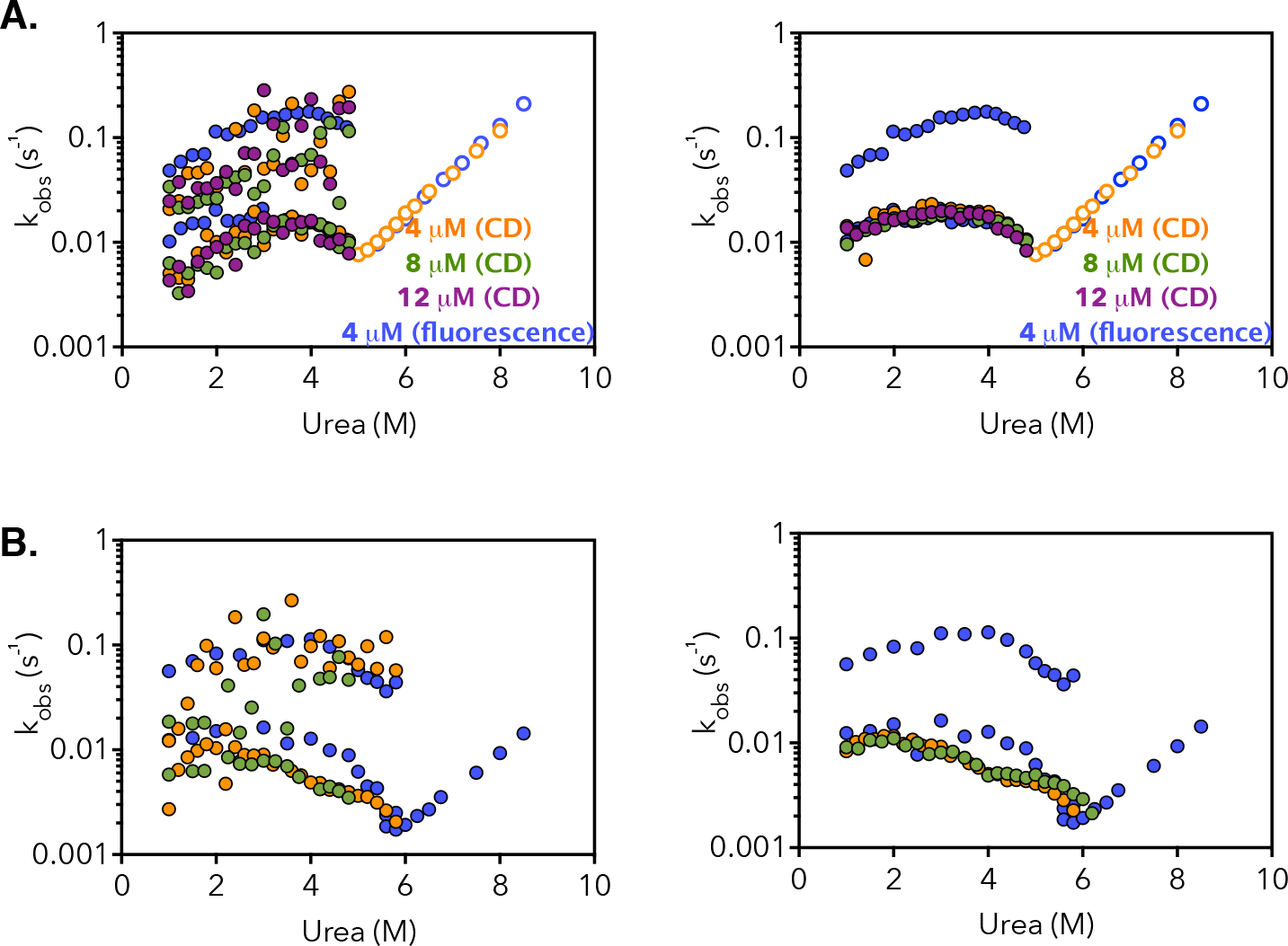
Refolding (solid circles) and unfolding (empty) rate constants of CheYHisF-2 (A) and G213I (B) obtained by fluorescence at 4 μM (blue) and CD at 4 μM, 8 μM and 12 μM. The CD refolding traces in were fit to either an equation describing biexponential decay (left) or to a monoexponential decay (right).

**Figure S9.**
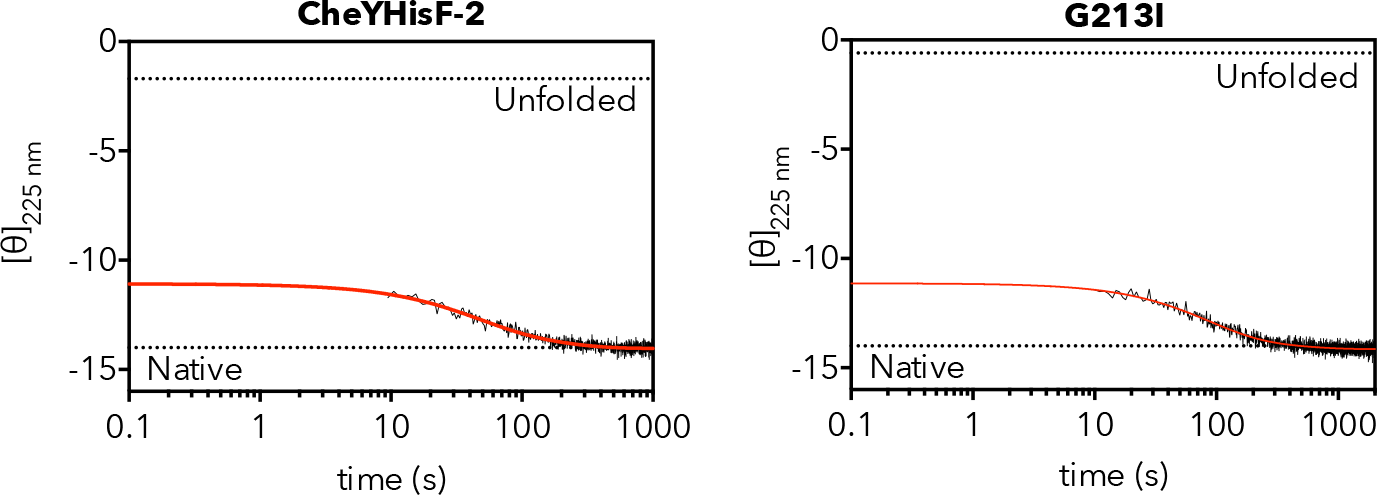
Refolding kinetic traces of CheYHisF-2 and G213I recorded using CD at 4 μM protein concentration. The dashed lines represent the signals of native or unfolded protein obtained from the fit of pre- and post-transition region of the equilibrium unfolding experiments. Fit of the traces to an equation describing biexponential decay is shown in red.

**Figure S10.**
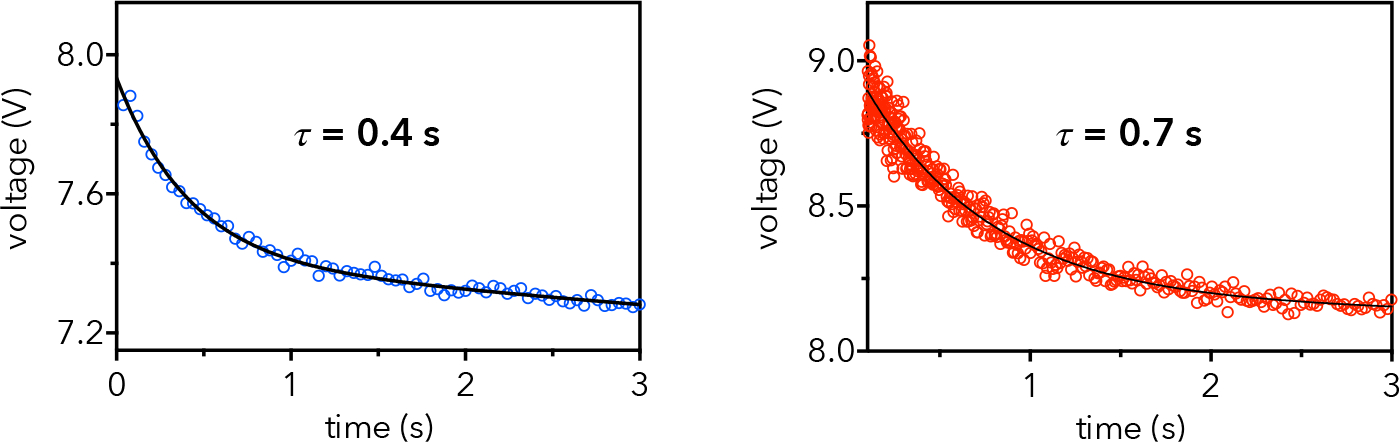
Kinetic unfolding trace of the intermediate I_ON._ CheYHisF-2 (blue) or G213 mutant (red) unfolded in 8.0 M urea was diluted 2-fold and allowed to refold for 18 s to populate I_ON_. The solution was then transferred to 8.5 M urea and the I_ON_ unfolding monitored by fluorescence (excitation = 280 nm, emission > 320 nm). The data was fit to a monoexponential decay plus a linear factor to account for decay of the intermediate and any changes due to protein accrued during the time delay.

## Notes

### Competing Interest Statement

The authors have declared no competing interest.

